# GDFold2: a fast and parallelizable protein folding environment with freely defined objective functions

**DOI:** 10.1101/2024.03.13.584741

**Authors:** Tianyu Mi, Nan Xiao, Haipeng Gong

## Abstract

An important step of mainstream protein structure prediction is to model the 3D protein structure based on the predicted 2D inter-residue geometric information. This folding step has been integrated into a unified neural network to allow end-to-end training in state-of-the-art methods like AlphaFold2, but is separately implemented using the Rosetta folding environment in some traditional methods like trRosetta. Despite the inferiority in prediction accuracy, the conventional approach allows for the sampling of various protein conformations compatible with the predicted geometric constraints, partially capturing the dynamic information. Here, we propose GDFold2, a novel protein folding environment, to address the limitations of Rosetta. On the one hand, GDFold2 is highly computationally efficient, capable of accomplishing multiple folding processes in parallel within the time scale of minutes for generic proteins. On the other hand, GDFold2 supports freely defined objective functions to fulfill diversified optimization requirements. Moreover, we propose a quality assessment (QA) model to provide reliable prediction on the quality of protein structures folded by GDFold2, thus substantially simplifying the selection of structural models. GDFold2 and the QA model could be combined to investigate the transition path between protein conformational states, and the online server is available at https://structpred.life.tsinghua.edu.cn/server_gdfold2.html.

## Introduction

Proteins are critical components of living organisms and provide the essential material basis for life processes. The function of a protein is typically dictated by its three-dimensional structure, as demonstrated by the general regulation mechanism of protein function and property through conformational change^1^. Hence, the acquisition of accurate three-dimensional structures of proteins is not only indispensable for mechanistic exploration in life sciences but also pivotal in advancing fields such as medicine and pharmacology. Unlike traditional protein structure determination that relies on costly and time-consuming biological experiments, the advancement of deep learning technologies has elicited unprecedented progress in the field of protein structure prediction, allowing accurate and fast inference of protein structures from their amino acid sequences, as witnessed by the critical assessment of protein structure prediction (CASP) competition^2^ in the past decade. In CASP12, RaptorX-Contact^3^ first applied the deep residual convolutional neural network to improve the prediction of inter-residue contacts from multiple sequence alignment (MSA), which effectively constrained the conformational sampling space in the downstream structural modeling process. AlphaFold1^4^ upgraded the contact prediction to distance prediction in CASP13, followed by trRosetta^5^ which proposed to predict the inter-residue orientations as additional geometric constraints. Distributions of these predicted geometric constraints could be converted to pseudo energies in protein folding environments like Rosetta^6^ to construct compatible protein structural models through gradient-based optimization. AlphaFold2^7^ in CASP14, followed by RoseTTAFold^8^, integrated the inter-residue geometry prediction and geometry-constrained structural modeling in a unified neural network, which markedly suppressed the prediction errors through end-to-end training, enhancing the precision of monomeric protein structure prediction to a level comparable to experimental resorts. Subsequent efforts included but not limited to open-sourcing the AlphaFold2 framework (OpenFold^9^ and UniFold^10^), prediction of protein-involved complex structures (AlphaFold-Multimer^11^, DeepMSA2^12^, RoseTTAFold All-Atom^13^, NeuralLPexer^14^ and AlphaFold3^15^), replacing the MSA processing by the protein language model to allow structure prediction from single sequences (ESM-Fold^16^, OmegaFold^17^, HelixFold-single^18^, RaptorX-Single^19^, RGN2^20^ and trRosettaX-Single^21^), as well as exploring multiple conformational states through MSA manipulation in AlphaFold2 (AF-Cluster^22^).

Despite the great success, AlphaFold has limited power in exploring dynamic protein structures. AF-Cluster partially overcomes this problem by iteratively feeding MSA sub-clusters to AlphaFold2, but still confronts the challenges of the arbitrariness and subjectiveness in MSA manipulation, randomness and uncertainty in the prediction results, as well as the high demand for computational resources. As a contrast, the Rosetta-based folding procedure of RoseTTAFold and trRosetta are, in principle, capable of sampling various conformations compatible with the predicted geometric constraints and thus capture the protein dynamics to some extent, despite the mildly declined prediction accuracy due to the isolation of constraint prediction and structural modeling. The Rosetta environment employed by the RoseTTAFold/trRosetta pipeline, however, has the following deficits: 1) the folding process is slow, frequently taking hours for a medium-sized polypeptide chain; 2) the folding framework is rigid in that all predicted geometric constraints have to be artificially converted into the style of statistical potentials. The second point, particularly, restricts its application in modeling protein structures using diversified constraints. For instance, in our recent work, we developed a lightweight protein structure prediction method SPIRED^23^, which utilizes 2D information to predict *L* (*i*.*e*. the number of residues) sets of C_*α*_ atom coordinates, each taking the local coordinate system of one individual residue as reference. Modeling the protein structures compatible with such prediction results is feasible in Rosetta only by converting the information-abundant C_*α*_ coordinates into distributions of inter-residue distances, a procedure that clearly leads to severe information loss. In another work, we developed a novel protein structure prediction framework Cerebra^24^, which significantly improves the computational efficiency by predicting multiple sets of atomic coordinates in parallel and leveraging their complementarity for prediction error suppression. In each set of prediction results, Cerebra outputs the relative translations and rotations (in quaternions) of the local coordinate systems of all residues in the reference frame of one anchor residue. Unfortunately, structural modeling given such prediction results is unsupported by the available energy definitions of the Rosetta environment. Therefore, the development of a more efficient protein folding environment with a higher degree of freedom in the objective function definition is still awaiting investigation.

In this work, we developed a novel protein folding environment, GDFold2 (<u>g</u>radient <u>d</u>escent based <u>fold</u>ing environment version <u>2</u>, distinguished from GDFold^25^), to fulfill the aforementioned demand. Firstly, it is capable of integrating various sources of predicted geometric information with statistical constraints to rapidly construct satisfactory allatom protein structures. Secondly, it supports highly flexible and fully customizable constraining loss functions, which can be rewritten by users according to their definitions and requirements. Here, we provide three predefined folding modes as references for users, which aim to generate all-atom protein structures compatible with the prediction results of SPIRED, Cerebra and RoseTTAFold/trRosetta, respectively. Moreover, GDFold2 is highly computationally efficient, faster than pyRosetta by at least one order of magnitude, and allows parallelized structural modeling. We also trained a quality assessment (QA) model to predict the ranking of quality for the structural decoys generated by multiple GDFold2 folding events. This scoring model can reliably assist users in selecting high-quality structural models that are compatible with their predicted constraints, and the combination of GDFold2 and the QA model enables the rapid identification of the transition path between different conformational states.

## Methods

As shown in Figure 1, the overall process of our method consists of two main stages. In the first stage, the geometric information predicted from various models (or converted from known structures) is converted into constraints, and then GDFold2 folds the protein structure based on these constraints. In the second stage, the multiple structural decoys produced by GDFold2 are evaluated by the QA model for quality ranking and model selection.

**Figure 1.**
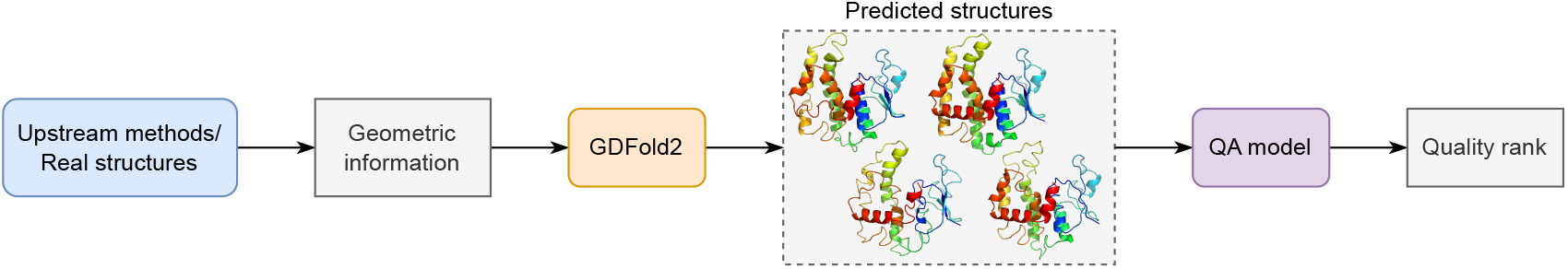
Overview of the entire method. Given the geometric information either predicted by upstream methods or converted from real structures, GDFold is capable of rapidly generating multiple structure decoys, which are further selected based on the evaluation of the QA model.

### Overview of the GDFold2 pipeline

The pipeline of GDFold2 is shown in Figure 2A. In this folding environment, 2D geometric information predicted from various methods or extracted from known structures (Figure 2B) and statistics summarized from high-resolution protein structures (Figure 2C) are converted into geometric constraints and statistical constraints, respectively, which are then taken as optimization aims for the user-defined loss functions in the neural network (named atomic coordinate network) for atomic coordinate generation. Specifically, coordinates of C_*α*_ atoms and relative rotations of the local coordinate systems of all residues are registered as learnable parameters of the atomic coordinate network, whose values are automatically optimized during the gradient-descent-based minimization of loss functions. Upon convergence, the coordinates of all atoms could be reconstructed from the optimized learnable parameters, based on prior knowledge of the intra-residue and peptide-bond internal coordinate information.

**Figure 2.**
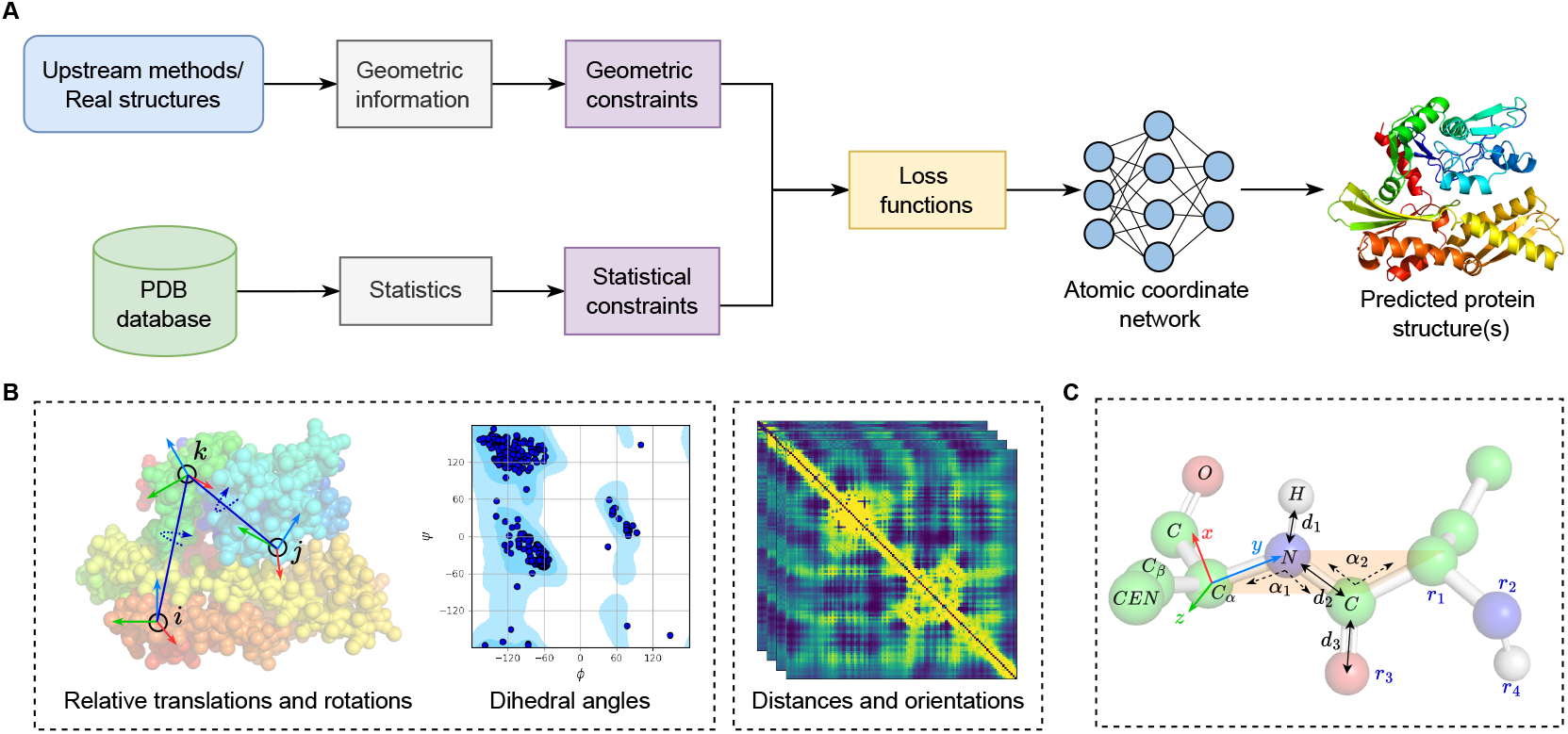
Schematic representation of the GDFold2 pipeline. **A)** Workflow of the GDFold2 pipeline. **B)** Various types of predicted geometric information are obtained from different models. The relative translations and/or rotations of residue pairs, backbone dihedral angles, as well as the probability distributions of inter-residue distances and orientational angles are shown as examples here, from left to right. **C)** Statistical parameters describing the general characteristics of polypeptide chains are obtained from high-resolution protein structures.

### Geometric constraints

Different protein structure prediction methods often output diversified geometric information as prediction results. In SPIRED and Cerebra, a set of *K* residues are selected as anchor residues and their local coordinate systems are established with the C_*α*_ atoms placed at origins (Figure 2B, left panel). Prediction results include the translational vectors of all residues within each reference frame of the *K* anchor residues as well as the main-chain dihedral angles. Cerebra predictions also include the residue rotations (in quaternions), which in combination with translations allow estimation of the positional vectors (**v**) of C, N and C_*β*_ atoms in addition to the C_*α*_ atoms within each of the *K* anchor reference frames. In the folding process of GDFold2, the same relative vectors could be computed from the learnable parameters, which thus allows evaluating the degree of consistency between protein structure under optimization and the prediction results, as shown in Equation 1:

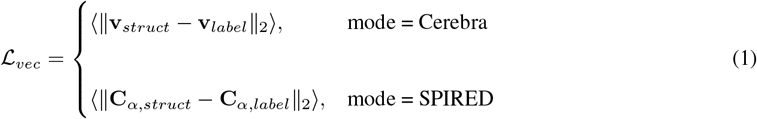

where ⟨∥ · ∥_2_⟩ denotes the *L*_2_ norms averaged over all residues and over all reference frames, while *struct* and *label* refer to the protein structure under optimization and the label converted from the predict results of SPIRED/Cerebra, respectively. The detailed implementation of ℒ_*vec*_ could be found in Algorithm 1 and Algorithm 2 (in **Supplementary Information 4**) for Cerebra and SPIRED, respectively. Moreover, main-chain dihedral angles estimated from the protein structure under optimization are also evaluated against prediction results in a similar manner (see Algorithms 3, 4 and 5 in **Supplementary Information 4** for details).

As for RoseTTAFold/trRosetta, the predicted probability distributions of inter-residue distances and orientational angles (Figure 2B, right panel) are converted into loss functions in the format of statistical potentials, following the “trFold.py” script of RoseTTAFold.

### Statistical constraints

Proteins should follow the chemical requirement of polypeptides during conformational changes. Hence, the universal characteristics of polypeptides summarized from high-resolution experimental structures could serve as statistical constraints. We analyzed protein structures with resolution < 3 Å in the CATH^26^ database and extracted the following statistical parameters: peptide bond length, distance between neighboring residues, angles between the adjacent covalent bonds within the peptide plane, van der Waals radii of backbone atoms, as well as vectors of carbonyl oxygen (O) and hydrogen (H) atoms in the peptide plane coordinate system and residue coordinate systems of common amino acids (Figure 2C, see **Supplementary Information 1.1** for details). Incidentally, to reduce the computational complexity, we substitute the side-chain of each amino acid residue with a centroid atom that has a larger van der Waals radius (see *CEN* in Figure 2C). Constraints constructed based on the above statistical data are applicable to all monomeric protein structure modeling. Detailed implementation of the construction of statistical constraints could be found in Algorithms 6, 7 and 8 (in **Supplementary Information 4**).

### Atomic coordinate network

Generally, protein folding refers to the physical process in which a protein transits from a random coil to a specific functional 3D structure. GDFold2 translates this process into the optimization of atomic coordinates of all amino acid residues based on predicted geometric information and statistical constraints. To achieve this goal, we establish a network with the C_*α*_ coordinates (in vectors) and the relative rotation matrices (in Euler angles) of all residues as learnable parameters, which are initialized randomly at the beginning. As mentioned previously, all constraining items are constructed in the form of loss functions required for updating neural network parameters, leveraging the reverse automatic differentiation mechanism of PyTorch^27^, which significantly enhances the computation speed. Convergence of the overall loss indicates that the current atomic coordinates obtained from the network conform to the predicted geometric features and statistical patterns of polypeptides. Moreover, the batch size of learnable parameters existing in the form of high-dimensional tensors corresponds to the number of predicted structures, allowing the network to parallelly update multiple sets of atomic coordinates, thereby greatly improving the prediction efficiency.

### 3D protein structure modeling

The 3D protein structure is built in the global Cartesian coordinate system. The local coordinate system of each amino acid residue is defined in the same way as AlphaFold2^7^, by placing the C_*α*_ atom at the origin, C atom on the *x*-axis and N atom in the *xy*-plane. Thus, given the optimized C_*α*_ coordinates and relative rotation matrices of all residues, the positional vectors of all atoms could be inferred in the global frame, based on prior knowledge about the intra-residue and peptide-bond internal coordinate information (see **Supplementary Information 1.1** for details). The detailed implementation could be found in Algorithms 9, 10, 11, 12 and 13 (in **Supplementary Information 4**).

The integral folding process of GDFold2 is shown in Algorithm 14 (in **Supplementary Information 4**). The structural models output by GDFold2 are further optimized by a modified pyRosetta-based FastRelax procedure, with the starting backbone coordinates applied as positional restraints (in terms of quadratic potentials). This relaxation process enables sampling of the complete side-chain (instead of the centroid atoms) as well as further improvement on the backbone.

### Overview of the quality assessment pipeline

As shown in Figure 3, the QA model adopts the graph neural network (GNN) architecture, which reads in node and edge features collected from an input protein structure and outputs the evaluation score for the structural quality. Details about the QA model are described in the following sections in four aspects: data preparation, feature design, network architecture and model training.

**Figure 3.**
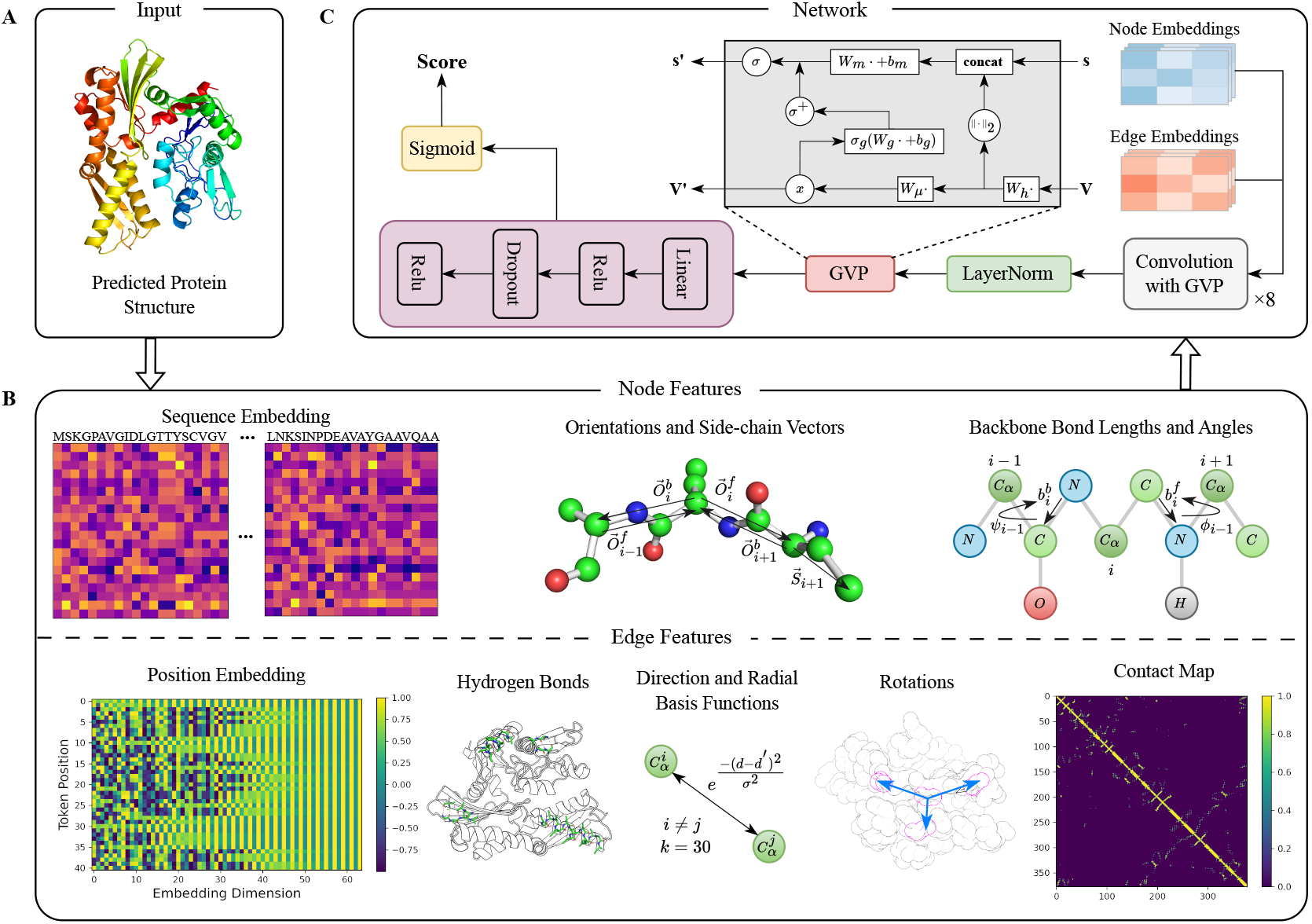
Schematic representation of the quality assessment pipeline. **A)** The protein structure under evaluation is taken as the raw input. **B)** Node and edge features are extracted from the input protein structure. Node features include sequence embedding, main-chain orientational and side-chain directional vectors as well as backbone bond lengths and angles. Edge features include positional embedding, one-hot encoding of hydrogen bonds, direction and distance between each pair of C_*α*_ atoms, C_*β*_ contact map as well as relative rotations between each residue and its top *K* spatially nearest neighboring residues. **C)** Network architecture of the QA model. The network embeds the entire features (in panel **B**) into node and edge embeddings and utilizes a graph-based message-passing mechanism to update and transfer the message between layers. A sigmoid activation function is connected with the last layer to output the evaluation score, which falls between 0 and 1.

### Data preparation

We randomly selected 4,169 and 5,452 target proteins from the training sets of SPIRED and Cerebra, respectively, and performed model inference to obtain the predicted geometric information. These targets range from 50 to 450 residues in length and have an average plDDT of 0.8735 as reported by SPIRED/Cerebra. To obtain decoy structures of varying predictive quality, we first designed the mask setting according to Equation 2:

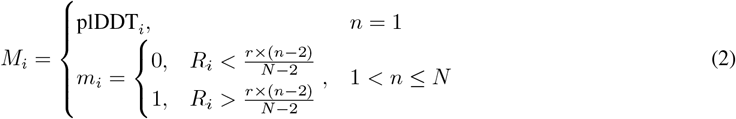

where *M*_*i*_ represents the masking weight for the *i*^*th*^ residue, *n* represents the *n*^*th*^ decoy structure, *r* and *N* are the upper limit of mask ratio and the number of decoy structures to be generated for each target, respectively, while *R* is a randomly generated float number between 0 and 1. Due to the distinct sensitivity to the masking weight, the setting of *r* is different according to the source of predicted geometric information (80% for Cerebra and 40% for SPIRED). *N* is determined by the number of anchor residues specified by Cerebra and SPIRED. These masks, acting as tensor weights, were then multiplied to the tensors of predicted constraints in the same shape of *N* × *L* to tune the scales of individual items. Hence, this procedure masked the predicted geometric information from loss calculation for randomly selected residues. Next, we used GDFold2 to fold these proteins parallelly, based on the masked geometric information. Finally, we obtained a total of 259,875 decoy structures for QA model training.

### Feature design

A protein structure is represented by a graph composed of nodes and edges in the QA model (Figure 3B). The node features characterize the information for each residue. Firstly, each residue of the protein is transformed into a 20-dimensional tensor to represent the sequence information. Then, in order to extract the topology features between residue pairs and characterize the overall conformation, we utilize the forward 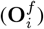 and backward orientations 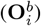 to represent the relationship between adjacent residues, and use a vector (**S**_*i*_) to describe the direction of side-chain centroid. Lastly, backbone bond lengths and angles are represented as scalar features. Specifically, peptide bond lengths are also embedded into forward 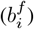 and backward 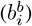 values to characterize the distance features between residue *r*_*i*_ and its succeeding residue *r*_*i*+1_ and preceding residue *r*_*i*−1_, respectively. Additionally, backbone dihedral angles *ϕ, ψ* and *ω* are embedded into tensors of (*L* − 1) × 2 dimension through sinusoidal and cosinusoidal trigonometric functions, respectively, to describe the relationship between adjacent planes.

The edge features in the graph are used to characterize the information between residue pairs. Here, we apply five edge features. Firstly, for each residue *i*, the top 30 spatially nearest neighboring residues are obtained by calculating the distance between their C_*α*_ atoms. The unit vector from residue *i* to its neighboring residue *j* is the direction feature, calculated as 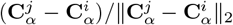. Then, we utilize the sinusoidal function and Gaussian radial basis function (RBF) to encode the direction and distance, respectively, to represent the relationship between these residue pairs. The embedding dimension of the direction feature and *σ* of the Gaussian RBF are hyper-parameters. In addition, hydrogen bonding is a kind of intra-molecular interaction that significantly contributes to the formation of protein secondary structures. Thus, we use the DSSP^28^ vacuum electrostatics model to encode the hydrogen bonds. Next, the relative rotations between each residue and its top 30 spatially nearest neighboring residues are calculated and then converted into the quaternion form to characterize the orientational relationship between residue pairs. Lastly, we use the contact map of C_*β*_ atoms with the contact cutoff of 8 Å to represent the overall structure information.

### Network architecture

The neural network takes as input the protein graph designed above and utilizes the message passing mechanism in Equation 3 and Equation 4: messages from neighboring nodes and edges are used to update node embeddings at each graph propagation step.

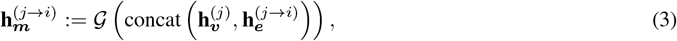

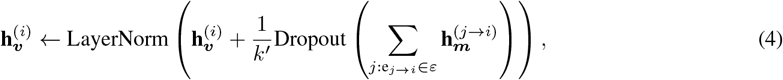

Here, 𝒢 is a sequence of graph vector perception (GVP)^29^ modules. The GVP module is designed to learn the vector-valued and scalar-valued functions for geometric vectors and scalars, respectively. In a GVP module, two linear projections are employed for the transformation of scalar and vector features. Before the transformation, scalar features are concatenated with the *L*_2_ norms of vector features, enabling the extraction of rotation-invariant information from vector features (Figure 3C, see Algorithm 15 in **Supplementary Information 4** for more details). 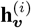 and 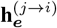 are the embeddings of the node *i* and directional edge (*j* → *i*), respectively, whereas 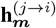 represents the message passing from node *j* to node *i. k*^*′*^ is equal to the number of incoming messages *k* unless the protein contains fewer than *k* amino acid residues.

### Model training

We randomly split 9,621 target proteins into training and validation sets with a ratio of 8:2. All decoy structures for one target protein are organized into one training mini-batch, and each training epoch involves traversing all target proteins. To balance the evaluation of global quality and local quality of predicted structures, we take the average value of lDDT and TM-score as the evaluation score and set the mean soft Spearman correlation coefficient^30^ between the predicted score and true evaluation value as the training objective (see following items for more details about evaluation metrics). Model training on an NVIDIA TITAN RTX 2080Ti GPU took 70 hours in total.

#### lDDT^31^

is a superposition-free score that indicates the difference in local inter-residue distances between the predicted structure and reference structure (Equation 5). In this study, we only calculate the lDDT score for C_*α*_ atoms.

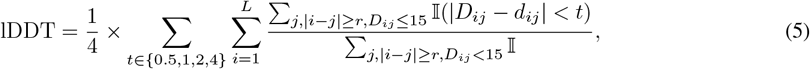

where I is the indicator function, *d* and *D* are the inter-residue distances in the predicted structure and reference structure, respectively, and *r* is the threshold for sequence separation.

#### TM-score^32^

is a metric used to assess the topological similarity between protein structures. According to Equation 6, TM-score ranges from 0 to 1, with the value of 1 indicating a perfect match between structures.

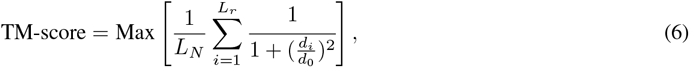

where *d*_*i*_ represents the distance between the *i*^*th*^ aligned residue pairs, *d*_0_ is a normalization scale, *L*_*N*_ denotes the original length of the protein, and *L*_*r*_ represents the number of aligned residues.

#### Score

is calculated from the average of TM-score and lDDT and is used as the evaluation metric of the QA model (Equation 7). It is not only sensitive to the local difference, but also indicates the degree of global match.

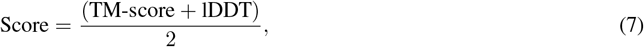

#### Spearman correlation coefficient^33^

(*ρ*) is commonly used to describe the strength of a monotonic relationship between two variables. According to Equation 8, Spearman correlation coefficient is calculated by utilizing the ranked values of both variables (*X, Y*).

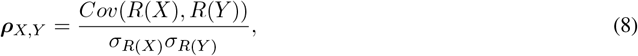

where *Cov* and *σ* denote the covariance and standard deviation, respectively, while *R* is the ranking operation.

## Results

### The folding process and performance of GDFold2

The folding process of GDFold2 can be divided into three main stages (Figure 4A), separated by the step points at which new loss terms are applied. The first stage optimizes the backbone structure of the target protein using the loss functions designed for the predicted geometric constraints. Due to the random initialization of learnable parameters in the atomic coordinate network, all atoms are located within a small sphere at the beginning. After the imposition of the predicted geometric constraints, the protein structure expands in dimension, gradually exhibiting features of the predicted geometry. In the second stage, multiple statistical constraint terms are employed as optimization objectives to fine-tune the local conformations, aiming to rectify the inconsistency between the predicted geometric information and the statistical regularities of polypeptides. After the first two stages, the protein backbone structure generally meets the prediction requirements. In the third stage, the van der Waals repulsion constraint is applied to eliminate the possible steric clashes between atom pairs. Eventually, a modified pyRosetta-based FastRelax procedure is engaged to sample the full side-chain conformations based on the coordinates of centroid dummy atoms (*i*.*e. CEN*) output by GDFold2. Besides the full sampling of side-chain rotamers, the main-chain hydrogen bond network and dihedral angles are also improved in this process. To accelerate the folding process, the learning rate is set to a large value (0.6) for the first stage but tuned to a small value (0.02) for the second and third stages. Incidentally, whenever a new constraint term is introduced as an optimization objective, the loss value rises up abruptly but anneals down rapidly, supporting their successful constraining effects.

**Figure 4.**
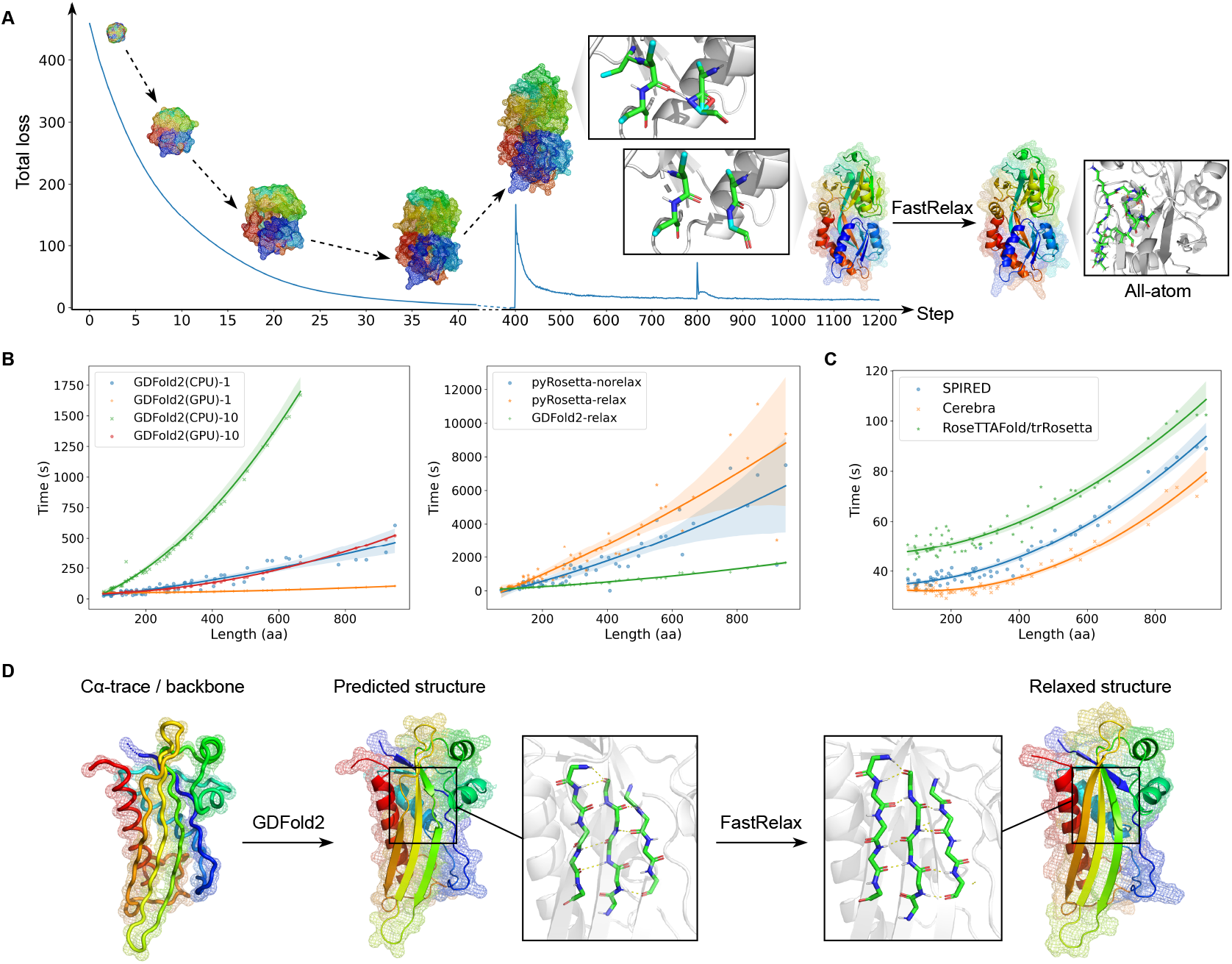
Folding process and performance of GDFold2. **A)** Evolution of the loss function and the protein structure in the folding process of GDFold2. The horizontal axis represents the number of optimization steps and the vertical axis represents the sum value of all applied losses. At steps 400 and 800, the statistical loss and van der Waals loss are added to the optimization process, respectively, where the insets present the gradually refined local backbone structures by these two loss functions. At the end of the folding process, typically at the step of 1200, a self-defined pyRosetta-based FastRelax process is applied to sample the full side-chain conformation, allowing for the construction of the all-atom structure model. **B)** Analysis of folding time for GDFold2. The analysis is performed by taking the SPIRED prediction as input, with the horizontal and vertical axes representing the protein length and the folding time, respectively. In the left subplot, the folding time of GDFold2 is evaluated on CPU and GPU with the batch size of 1 and 10, respectively, without the FastRelax process. In the right subplot, the folding time of the full GDFold2 pipeline (with the FastRelax process) is compared with Rosetta with and without structure relaxation. **C)** Comparison of the GDFold2 folding time when the geometric constraints predicted by SPIRED, Cerebra and RoseTTAFold/trRosetta are taken as input. **D)** Illustration of the role of the self-defined pyRosetta-based FastRelax process. The initial C_*α*_-trace/backbone predicted by SPIRED/Cerebra could be rapidly refined by GDFold2 to restore most of the backbone secondary structures. In the subsequent FastRelax process, the initial backbone coordinates of the GDFold2-folded proteins are positionally constrained, aiming for sampling the side-chain atoms and further optimizing the main-chain hydrogen networks without compromising the high prediction accuracies of SPIRED/Cerebra models.

We evaluated the folding time of GDFold2 on 65 randomly selected protein targets with length ranging from 73 to 949 residues, using the SPIRED predictions as geometric constraints (overall folding step = 1200; batch size = 1). According to the evaluation curve (Figure 4B, left), GDFold2 accomplishes folding within 1 minute on GPU for proteins of less than 400 residues. Even for large proteins of ∼1000 residues, the full folding process only takes 2 minutes, significantly faster than the traditional Rosetta-based folding (Figure 4B, right). Subsequently, in order to quantitatively compare the running time of GDFold2 and pyRosetta, we took the RoseTTAFold-predicted distributions of inter-residue distances and orientational angles of the selected protein targets as input, and employed GDFold2 (overall folding step = 1200; batch size = 10) and pyRosetta (default parameters from RoseTTAFold) to build the structural models, respectively. At the same level of quality for the generated structural models (see **Supplementary Information 1.2** for the case study), GDFold2 is markedly more efficient than pyRosetta: the per-structure average folding time of GDFold2 is far less than that of pyRosetta. Notably, excluding the MSA preparation time, the pure inference of geometric constraints by the upstream methods like SPIRED, Cerebra and RoseTTAFold/trRosetta typically fall into the time scale of seconds, faster than the downstream Rosetta-based structural modeling/refinement process by orders of magnitude. As a contrast, GDFold2 remarkably accelerates the folding process, rescuing this rate-limiting step back to the time scale of tens of seconds, no matter whether using the constraint styles of SPIRED, Cerebra or RoseTTAFold/trRosetta (Figure 4C).

The raw prediction results of SPIRED contain coordinate information for C_*α*_ atoms only. However, by converting the C_*α*_ coordinates into geometric constraints, GDFold2 is capable of replenishing the missing atoms. As shown by the example in Figure 4A&D, the all-atom model provided by GDFold2 demonstrates abundant structural details, *e*.*g*., secondary structure contents and side-chain conformations, while the nearly negligible decline in C_*α*_ TM-score (or rise in root-mean-square-deviation (RMSD)) supports the compatibility of this structural model with the raw SPIRED prediction. Hence, the non-statistical-potential-style loss function in GDFold2 successfully complements the 2D geometric prediction results by statistics derived from high-resolution protein structures, nearly without causing information loss. More GDFold2 folding cases could be found in the Supplementary Information of our SPIRED paper^23^. Unlike SPIRED, Cerebra prediction results involve full information required for all-atom structural reconstruction. However, GDFold2 folding based on Cerebra-type constraints can improve the local structural quality due to the inclusion of the statistical constraints in the optimization process (see **Supplementary Information 1.3** for details).

In the third stage of GDFold2 folding process, the van der Waals repulsion constraint employed for clash elimination is inclined to push the side-chain centroid dummy atoms of large residues farther away, thus disrupting the potential hydrogen bonds and secondary structure contents (Figure 4D, see the structure in the left inset). The pyRosetta-based FastRelax procedure can effectively fix this problem, but frequently elicits large deviations in the positions of backbone atoms, leading to considerable inconsistency between the relaxed protein structures and SPIRED/Cerebra predictions. To address this trouble, we implemented a specialized pyRosetta-based FastRelax procedure for GDFold2, by taking the backbone atom coordinates of GDFold2-folded structures as positional restraints during the simulated annealing processes of FastRelax. As shown in Figure 4D, the self-defined FastRelax method optimizes both the side-chain conformations and the secondary structure contents of the GDFold2-folded structure (see the structure in the right inset), while maintaining the high predictive accuracy of the main-chain structure.

Besides the conventional structure modeling for protein monomers, GDFold2 is capable of modeling the all-atom structures of protein complexes. Here, we take a dimeric glutathione transferase (PDB ID: 1LBK) as an example. For this protein complex, Cerebra could be engaged to provide the crude backbone structural model. In the structural modeling/refinement for this target, the all-atom structure model could be quickly generated, with a slight modification on the statistical loss terms to reinforce the lack of amino acid linkage across subunits (see **Supplementary Information 1.4** for details).

### Performance of the QA model in ranking and selection

We chose a number of proteins from the training sets of SPIRED and Cerebra, and employed the highly efficient GDFold2 pipeline to generate numerous decoy structures with a considerable degree of structural variation for each protein target by randomly masking the SPIRED and Cerebra predictions in the geometric loss function (Figure 5A). We then trained the QA model to learn the relationship between the decoy features and their structural quality. During the evaluation of our QA model, in order to avoid information leaking, we thus chose proteins in the CAMEO testing dateset^23^ that are non-homologous to those adopted for model training (with the maximal pairwise sequence similarity < 30%), selected RoseTTAFold predictions (rather than SPIRED/Cerebra ones used for model training) as constraints, and generated 24 structural models by GDFold2 for each protein target. Generally, the structural models produced for each individual protein target present a certain degree of structural dissimilarity, with an average deviation (*i*.*e*. max - min) of 0.0714 in the evaluation score (deviation of 0.048 in TM-score and 0.1016 in lDDT), since the RoseTTAFold prediction results (*i*.*e*. distributions of inter-residue distances and orientational angles) can only impose weak constraining effects. Here, we use the coefficient of variation (*c*_*v*_) to measure the degree of variation among the structural models generated by GDFold2:

**Figure 5.**
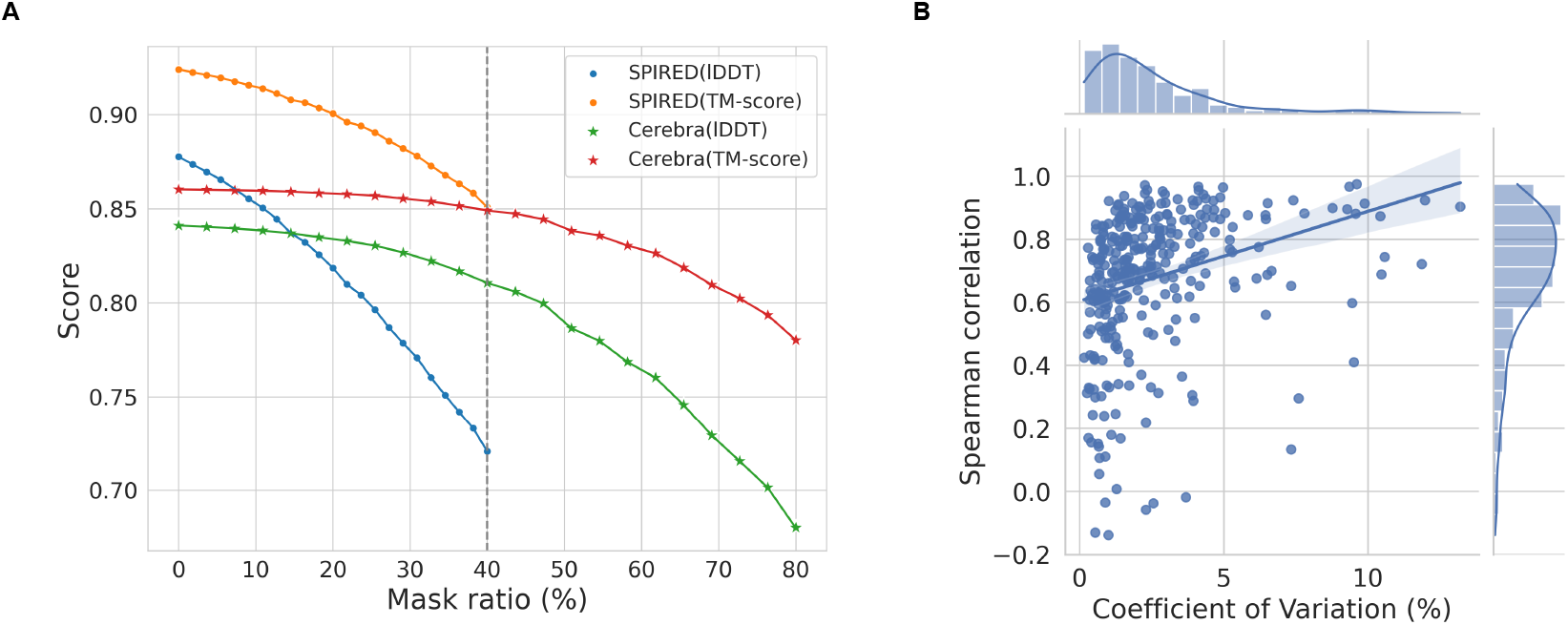
The QA model and its performance in ranking and selection. **A)** The effect of masking ratio on the variability of folded structures. The horizontal axis represents the mask ratio enforced on the geometric constraints, while the vertical axis represents the Score (*i*.*e*. TM-score and lDDT to the experimental structure) of generated structure decoys. Based on these results, the maximal masking ratio is set as 40% for SPIRED and 80% for Cerebra. **B)** Analysis on the structure decoys produced by GDFold2 based on RoseTTAFold-type constraints. Each dot in the plot denotes a protein. The horizontal axis represents the coefficient of variation (*c*_*v*_) of the evaluation score for all GDFold2-folded protein decoys, while the vertical axis represents the Spearman correlation coefficient (*ρ*) between the Score predicted by the QA model and the true evaluation score.

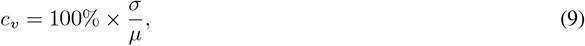

where *σ* and *µ* are the standard deviation and the mean of the true evaluation scores of the structural models, respectively. The total of 329 CAMEO testing protein targets present an average *c*_*v*_ of 2.54%, with 89.4% of the targets showing *c*_*v*_ values of < 5% (Figure 5B), reinforcing the small variation among the generated structural models. However, the QA model shows a strong capability of discerning these slightly different structural models: the Spearman correlation coefficient between the predicted Score and true evaluation score exceeds 0.6 for 75.1% of the testing targets, attaining an average value of 0.6767 for all targets. Expectedly, our QA model is more powerful at predicting the ranking of quality for structural models with larger degree of structural variations (Figure 5B).

Unlike the RoseTTAFold/trRosetta outputs, the prediction results of SPIRED and Cerebra are composed of atomic coordinate information, which intrinsically exerts strong constraining effects during GDFold2 folding. Hence, in the absence of masking, the structural models generated from the same set of SPIRED/Cerebra predictions usually show a high degree of structural similarity (see **Supplementary Information 2.1** for the effects of masking). In practical structural prediction, however, multiple model checkpoints with different random seeds are frequently collected as an ensemble to further enhance the prediction accuracy. Consequently, a good QA model should be able to select highquality structural models from the ensemble of GDFold2-folded structures using constraints predicted from various combinations of checkpoints and random seeds. Here, we evaluated our QA model in this aspect on 76 CAMEO protein targets. Specifically, we randomly selected 20 combinations of seeds and checkpoints (4 random seeds × 5 checkpoints) for Cerebra, fed the same MSA inputs to these Cerebra models to make prediction, and then employed GDFold2 to fold proteins using each individual set of prediction results as geometric constraints. Thus, for each of the 76 protein targets, we generated 20 structural models, corresponding to various combinations of seeds and checkpoints. As shown in Table 1, the best structures selected by the QA model outperform those chosen based on the plDDT reported by SPIRED/Cerebra, in terms of both the average TM-score and the average lDDT. Moreover, the QA model exhibits a stronger power at ranking the truly optimal structure as the best and among the top 3. More detailed results could be found in **Supplementary Information 2.2**.

**Table 1.** Comparison of the structural model selection by the QA model and the Cerebra-reported plDDT.

| Method | TM-score | IDDT | Truly Best as the Best | Truly Best in Top-3 |
| --- | --- | --- | --- | --- |
| **pIDDT** | 0.7153 | 0.7264 | 4/76 | 11/76 |
| **QA model** | 0.7269 | 0.7418 | 11/76 | 28/76 |

### Exploring the protein structural dynamics

The GDFold2 folding environment and the QA model developed in this work could be further integrated to investigate the structural dynamics of proteins. Here, we took the adenosine kinase (ADK) as an example. We collected two experimentally solved structures of this protein with highly distinct conformations (Figure 6A), *i*.*e*. the closed state (PDB ID: 1AKE) and the open state (PDB ID: 4AKE), and constructed a transition path between these two states following the procedure in Figure 6B. Specifically, we first converted these two structures into the Cerebra-style geometric constraints (*i*.*e*. relative translations and rotations between the local frames of amino acid residues). Next, we combined these two sources of geometric constraints into an overall constraint, while randomly masking the individual terms of various amino acid residues during GDFold2 folding (see Algorithm 16 in **Supplementary Information 4** for detailed implementation). This approach allows for rapid production of multiple structure decoys with highly diversified degrees of agreement with the hybrid geometric constraint. As shown in Figure 6C, the 50 generated structure decoys compose a path that reflects the gradual structural transition between the two terminal structural states both in the principal component analysis (PCA) and through visual inspection. We then clustered all structure decoys into 10 groups, chose a representative structural image with the highest quality from each cluster using the QA model, and further optimized their local structures by the self-defined pyRosetta-based FastRelax. Clearly, these structure images capture the continual structural transition process (Figure 6D). In Figure 6E, we evaluated the Rosetta energy and the QA-model score for the structural images as well as the two terminal structures. Nearly all structural images along the transition path show reasonably low Rosetta energy, supporting the physical plausibility of the identified path. Interestingly, at least in this case, the QA scores exhibit an evident negative correlation with the Rosetta energy values (Pearson correlation coefficient = -0.403). Hence, although energy terms are completely absent in the model training, our QA model tends to assign higher ranking scores to structures with lower potential energies, thus grasping the essential characteristics of protein structures.

**Figure 6.**
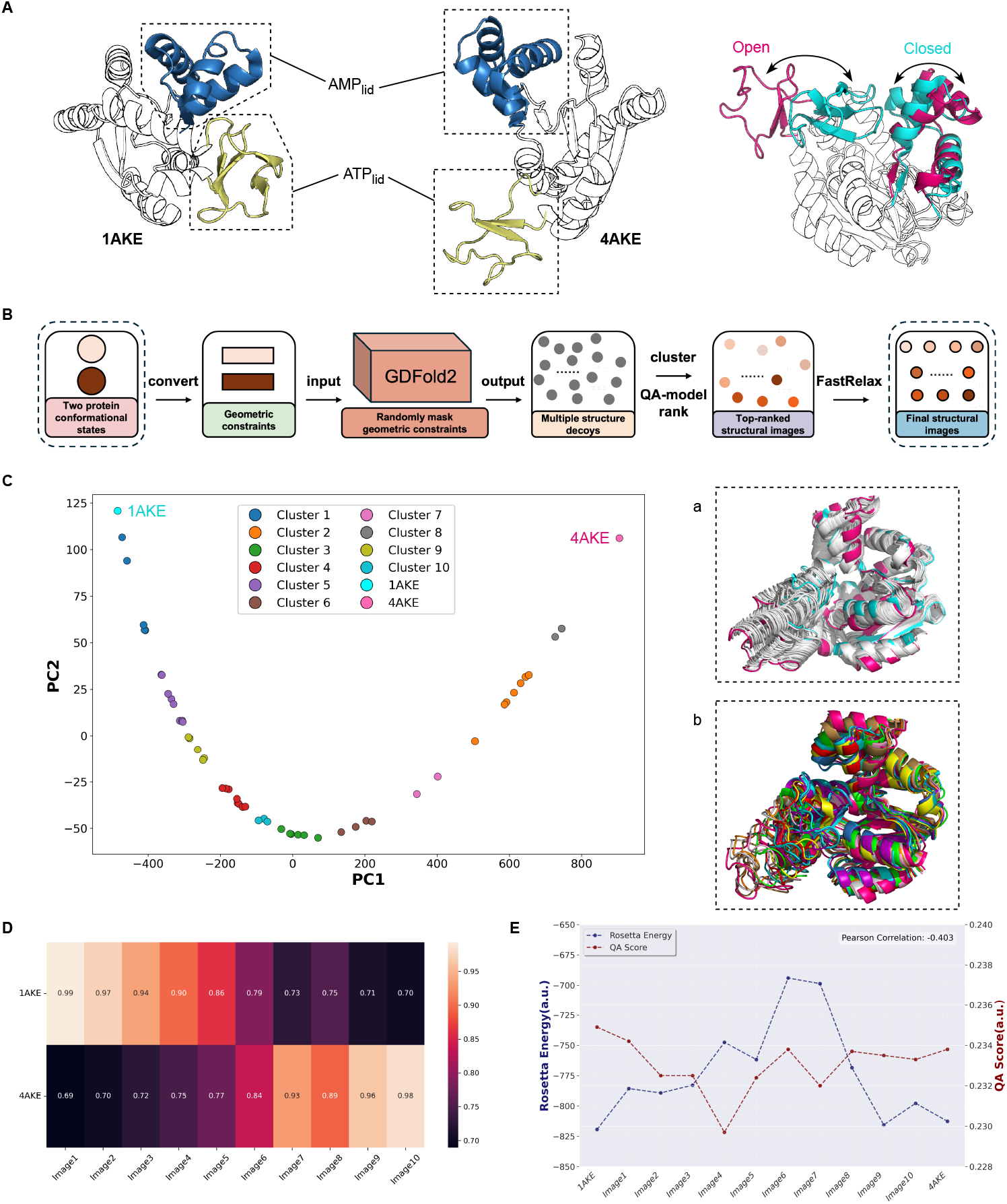
Transition path between two conformational states. **A)** Structural illustration of the two distinct conformational states of ADK. The AMP_lid_ (blue) and ATP_lid_ (yellow) present remarkable structural changes between the closed and the open states. **B)** Schematic illustration of the procedure for finding intermediate structural images between the two conformational states of a protein. **C)** The multiple structure decoys generated by GDFold2 through random masking compose a gradual transition path in the PCA analysis, with structures in the same cluster labeled in the same color. In the right subplot (**a**), all structural decoys (gray) are superposed onto structures of the closed (cyan) and open (hotpink) states. In the right subplot (**b**), representative structure images of all clusters are superposed and colored in the same style as in the PCA analysis. **D)** TM-scores between the representative structure images and the two PDB structures. **E)** Rosetta energy score and the QA-model score are evaluated on representative structure images along the transition path between the two conformational states. The structure images are further optimized by the FastRelax process for side-chain sampling. Notably, the magnitude of the QA score is unimportant since the objective functions in the QA model training is designed for evaluating the relative ranking between structure decoys only.

Nevertheless, in this exemplar investigation, we showcase the potential of the GDFold pipeline in the rapid generation of a physically plausible transition path linking two structural states for a protein. In practice, when the experimental structures are absent, AlphaFold2 could be used to produce multiple conformational states through MSA manipulation. In **Supplementary Information 3.1**, we show the generation of two distinctive structure states for ADK using different MSA clusters as inputs for AlphaFold2. With these two structural models as the source of geometric constraints, the transition path could be created quickly following the same procedure.

Besides the transition path, GDFold2 and the QA model could also be integrated to sample possible intervening states when more than two known structural states are taken as inputs. The structure decoys created by this means typically present low Rosetta energy. Hence, multiple short molecular dynamics trajectories could be run from these generated decoys in a parallelized manner, from which the Markov state model (MSM)^34^ could be constructed to analyze the kinetic behavior of the target protein given the input structural states.

## Conclusion

In this study, we have developed a novel protein folding environment, GDFold2, which allows freely defined objective functions for protein structure generation. Loss functions have been designed as examples for three application scenarios, with highly diversified geometric information predicted from various prediction methods. In all scenarios, the predefined loss function can effectively constrain the protein folding process and prompt the production of high-quality protein structural models in combination with statistical constraints derived from high-resolution protein structures. The overall framework is customizable, allowing users to define their own loss functions for fulfilling specific practical requirements. In comparison with popular protein folding environments like Rosetta, GDFold2 exhibits advantages in the significantly reduced folding time, the availability of parallelized folding and the high degree of freedom in the definition of objective functions. Additionally, a QA model has also been developed for reliably estimating the quality ranking of protein structures folded by GDFold2, which further simplifies the model selection process. Integration of GDFold2 and the QA model allows rapid exploration of the protein structural dynamics, particularly the transition path between different conformational states.

## Data availability

All data are available at https://zenodo.org/doi/10.5281/zenodo.10809439.

## Code availability

The codes of GDFold2 are available at https://github.com/Gonglab-THU/GDFold2. An online server is available at https://structpred.life.tsinghua.edu.cn/server_gdfold2.html.

## Author contribution

H. G. proposed the concept and theory. For the GDFold2 and QA models, T. M. and H. G. proposed the initial model design, while T. M. implemented coding, model training and testing. T. M. and N. X. conducted the data analysis. T. M., N. X. and H. G. wrote the manuscript. All authors agreed with the final manuscript.

## Acknowledgments

This work has been supported by the Ministry of Science and Technology of China (#2023YFF1204400), by the National Natural Science Foundation of China (#32171243) and by the Beijing Frontier Research Center for Biological Structure.

## Ethics declarations

### Competing interests

The authors declare no competing interests.

## Supplementary Information

### 1 Supplementary Details for GDFold2

#### 1.1 Statistical parameters used for the construction of statistical constraints

**Table S1.** Statistical values for scalars and vectors.

| Item | Value |
| --- | --- |
| $l$ | 1.33 |
| $d$ | 3.80 |
| $\theta_a$ | 0.520 |
| $\theta_b$ | 0.449 |
| $r_{C_\alpha}, r_C, r_{C_\beta}, r_{CEN}$ | 1.70 |
| $r_N$ | 1.55 |
| $r_O$ | 1.52 |
| $r_H$ | 1.20 |
| $\mathbf{v}_O$ | [-0.6653, -1.0348, 0.0028] |
| $\mathbf{v}_H$ | [-0.4889, -0.8837, 0.0] |

**Table S2.** Statistical internal coordinates of backbone atoms.

| Amino acid | C | N | $C_\beta$ | $CEN$ |
| --- | --- | --- | --- | --- |
| GLY | [1.516, 0, 0] | [-0.574, 1.335, 0] | NA | NA |
| ALA | [1.524, 0, 0] | [-0.525, 1.361, 0] | [-0.533, -0.775, -1.196] | [-0.532, -0.775, -1.195] |
| CYS | [1.522, 0, 0] | [-0.521, 1.361, 0] | [-0.540, -0.570, -1.312] | [-1.268, -1.274, -1.474] |
| GLU | [1.524, 0, 0] | [-0.527, 1.360, 0] | [-0.525, -0.778, -1.208] | [-1.623, -1.894, -1.882] |
| ASP | [1.526, 0, 0] | [-0.524, 1.361, 0] | [-0.526, -0.779, -1.208] | [-1.258, -1.443, -1.455] |
| PHE | [1.523, 0, 0] | [-0.516, 1.363, 0] | [-0.528, -0.775, -1.208] | [-1.681, -1.818, -1.539] |
| ILE | [1.525, 0, 0] | [-0.492, 1.373, 0] | [-0.514, -0.795, -1.214] | [-1.331, -1.404, -1.601] |
| HIS | [1.522, 0, 0] | [-0.525, 1.360, 0] | [-0.523, -0.784, -1.208] | [-1.581, -1.688, -1.511] |
| LYS | [1.524, 0, 0] | [-0.524, 1.361, 0] | [-0.526, -0.778, -1.207] | [-1.849, -2.234, -1.984] |
| MET | [1.523, 0, 0] | [-0.523, 1.362, 0] | [-0.529, -0.771, -1.208] | [-1.620, -2.064, -1.667] |
| LEU | [1.523, 0, 0] | [-0.520, 1.363, 0] | [-0.535, -0.774, -1.211] | [-1.619, -1.857, -1.363] |
| ASN | [1.524, 0, 0] | [-0.535, 1.356, 0] | [-0.523, -0.800, -1.179] | [-1.318, -1.449, -1.392] |
| GLN | [1.524, 0, 0] | [-0.525, 1.361, 0] | [-0.517, -0.775, -1.215] | [-1.690, -1.947, -1.757] |
| PRO | [1.524, 0, 0] | [-0.568, 1.350, 0] | [-0.568, -0.637, -1.279] | [-1.296, 0.973, -1.528] |
| SER | [1.523, 0, 0] | [-0.530, 1.358, 0] | [-0.527, -0.765, -1.198] | [-0.698, -0.859, -1.706] |
| ARG | [1.524, 0, 0] | [-0.523, 1.362, 0] | [-0.558, -0.831, -1.146] | [-1.988, -2.416, -2.408] |
| THR | [1.524, 0, 0] | [-0.517, 1.363, 0] | [-0.506, -0.799, -1.215] | [-0.937, -1.026, -1.537] |
| TRP | [1.523, 0, 0] | [-0.522, 1.361, 0] | [-0.525, -0.774, -1.210] | [-1.500, -2.164, -1.648] |
| VAL | [1.524, 0, 0] | [-0.493, 1.373, 0] | [-0.513, -0.795, -1.215] | [-0.937, -1.304, -1.406] |
| TYR | [1.523, 0, 0] | [-0.519, 1.362, 0] | [-0.526, -0.776, -1.209] | [-1.757, -1.952, -1.625] |

As described in **Methods**, atomic-level parameters summarized from high-resolution protein structures are used to construct statistical constraints. The parameters include peptide bond length (*l*), the distance between adjacent residues (*d*), angles between adjacent covalent bonds within the peptide plane (*θ*_*a*_ and *θ*_*b*_), van der Waals radii of backbone atoms 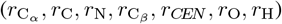, as well as the vectors of O and H atoms in the peptide plane coordinate system (**v**_O_ and **v**_H_), as listed in Table S1. The internal coordinates of backbone atoms of 20 common amino acids (Table S2) are calculated in the local coordinate system of each residue, taking C_*α*_ as the origin ([0, 0, 0]).

#### 1.2 GDFold2 is significantly more efficient in folding than pyRosetta

**Figure S1.**
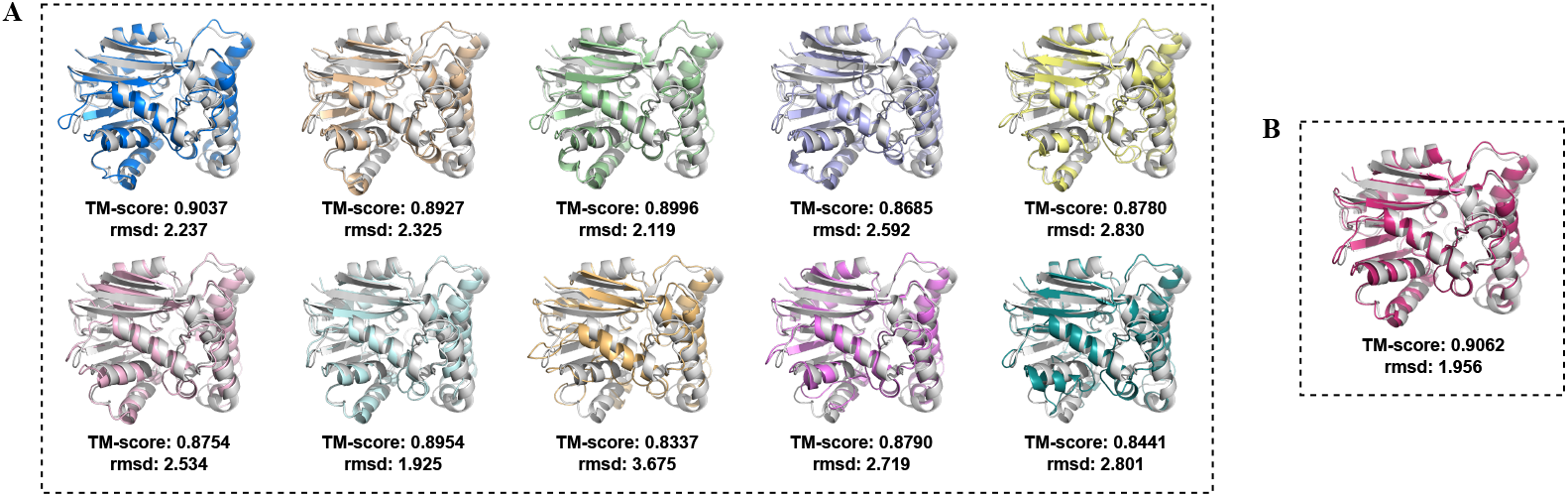
Comparison of the protein structures folded by GDFold2 and pyRosetta. **A)** Multiple prediction results output by GDFold2 through parallel folding (batch size = 10). **B)** Single prediction result output by pyRosetta. All of the generated structures are superimposed into the native structure (colored gray).

As shown in Figure S1A, multiple predicted protein structures are parallelly folded by GDFold2 using the RoseTTAFold predictions as constraints and are then relaxed following our modified pyRosetta-based FastRelax procedure. The total runtime to obtain all these prediction results is just 756.36 seconds. The single structure folded by pyRosetta^1^ with FastRelax is generated by the same RoseTTAFold-predicted constraints (Figure S1B), which consumes 2614.89 seconds. The structural models generated by the two methods present the same level of quality. However, the folding efficiency of GDFold2 is much higher than that of the pyRosetta environment.

#### 1.3 GDFold2 folding improves the local structural quality for Cerebra predictions

**Figure S2.**
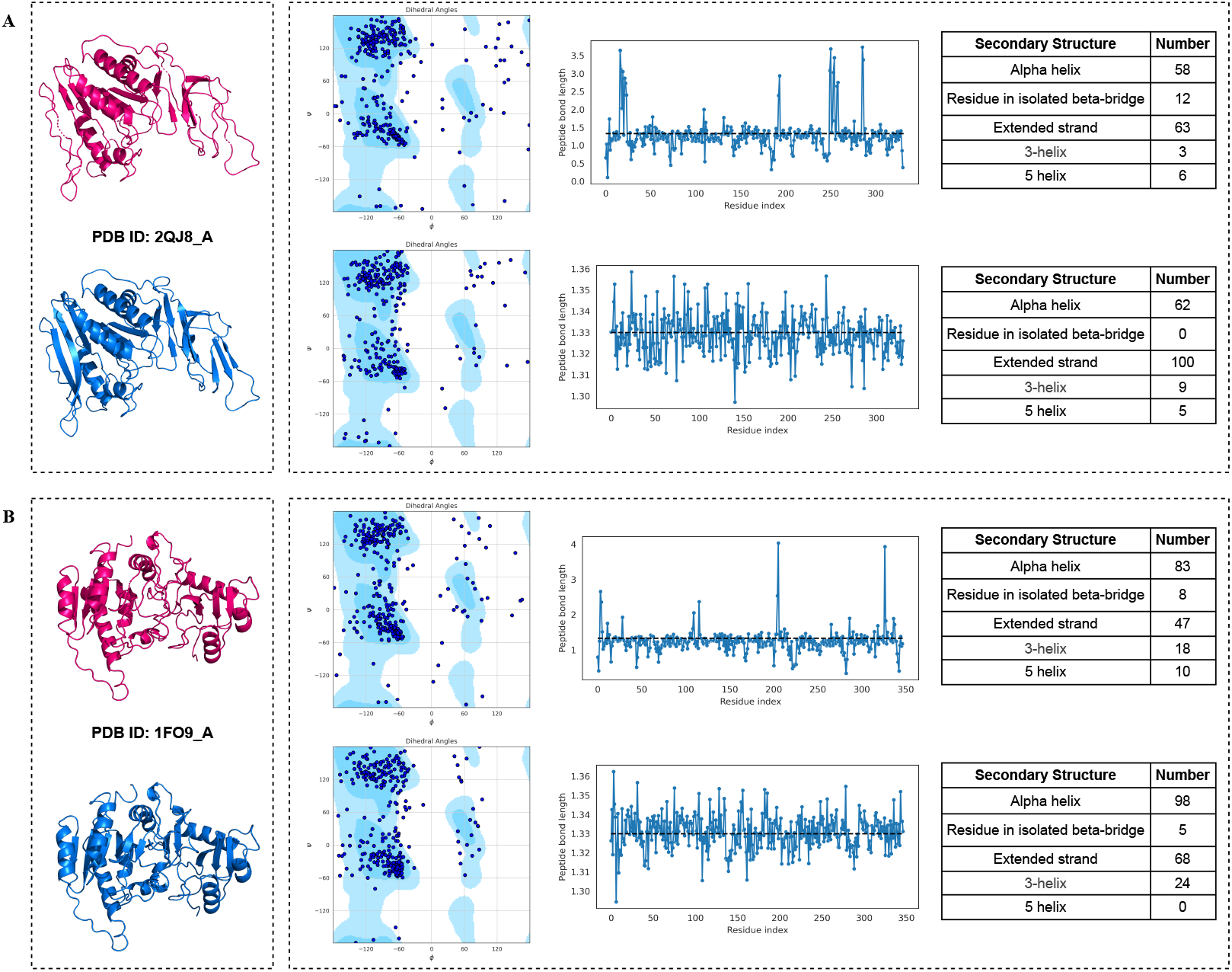
Comparison of structures generated from by GDFold2 and by direct construction based on Cerebra predictions. The comparison is shown for two cases: **A)** PDB ID: 2QJ8, chain A; **B)** PDB ID: 1FO9, chain A. In both cases, the structures directly output from Cerebra (red) and their properties are shown in the upper row, whereas the GDFold2-folded structures (blue) and their properties are shown in the lower row.

As shown in Figure S2, the incorrectly predicted local geometric information may lead to the appearance of local chain breakage at the peptide bonds (unrealistic distance between the adjacent C and N atoms), unreasonable distribution of main-chain torsion angles (violation of the Ramachandran plot) and damage on secondary structures in the raw Cerebra predictions. The protein structures generated by GDFold2 using Cerebra-predicted information and statistical constraints exhibit refinement in the local structural properties including the Ramachandran plot of main-chain torsion angles, the distribution of peptide-bond lengths as well as the number of residues in secondary structure contents (especially for categories of isolated beta-bridge and extended strand) based on DSSP^2^ calculation.

#### 1.4 GDFold2 can model the protein complex structure

**Figure S3.**
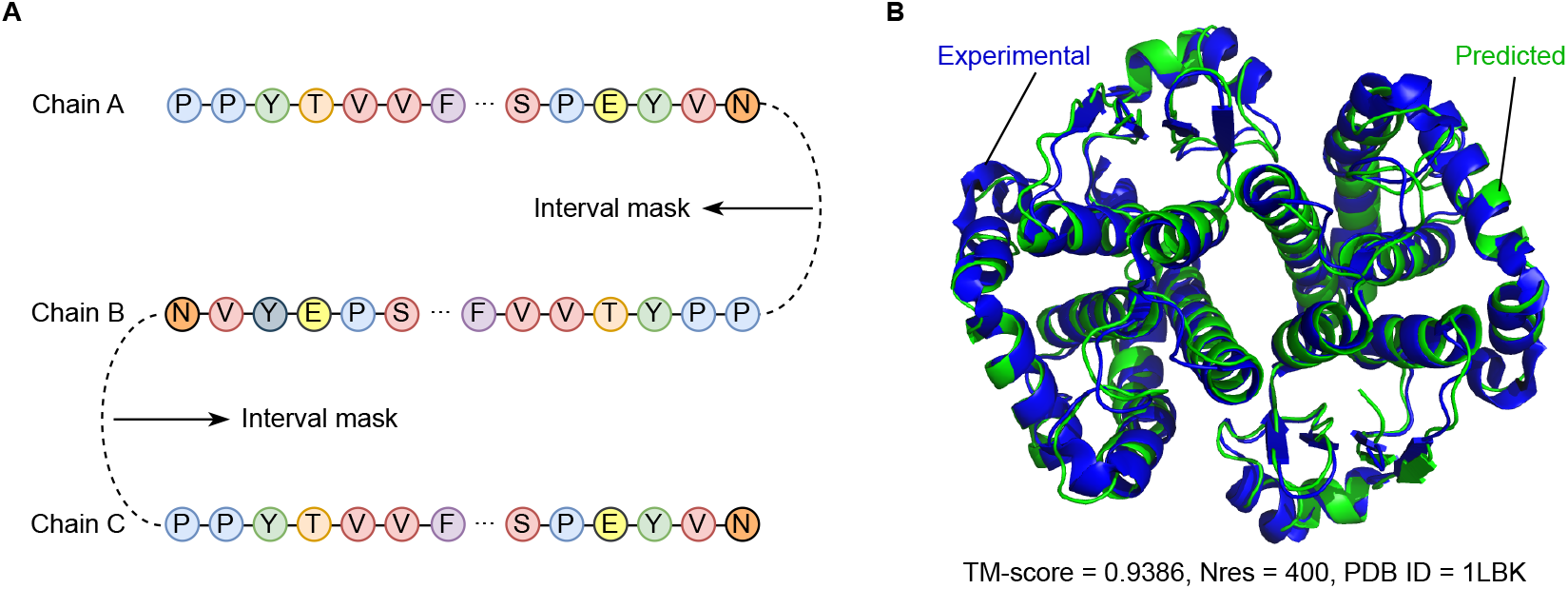
Folding the protein complex. **A)** A schematic illustration for modeling the protein complex structure by GDFold2. The statistical loss terms enforcing the peptide bonding between adjacent residues should be masked when the sequence passes the boundary of different polypeptide chains. **B)** The structure refolded by GDFold2 (green) is superposed to the experimental structure (blue).

Here, we show how to use GDFold2 to model the structures of protein complexes. In the conventional GDFold2, all amino acids are sequentially fed to GDFold2, and the statistical loss is automatically applied to adjacent residues to enforce the formation of proper peptide bond. In the modeling of protein complex structures, these statistical loss terms should be masked off when the adjacent residues cross the boundary of two subunits (Figure S3A). In Figure S3B, we present the folding result for a dimeric glutathione transferase (PDB ID: 1LBK). The geometric information was first predicted for this protein complex using Cerebra. Then GDFold2 was used to generate the all-atom structure model. The final structure resembles the experimental structure, with the TM-score exceeding 0.93.

### 2 Supplementary Details for the QA Model

#### 2.1 Case study for the decoy structures folded by GDFold2

**Figure S4.**
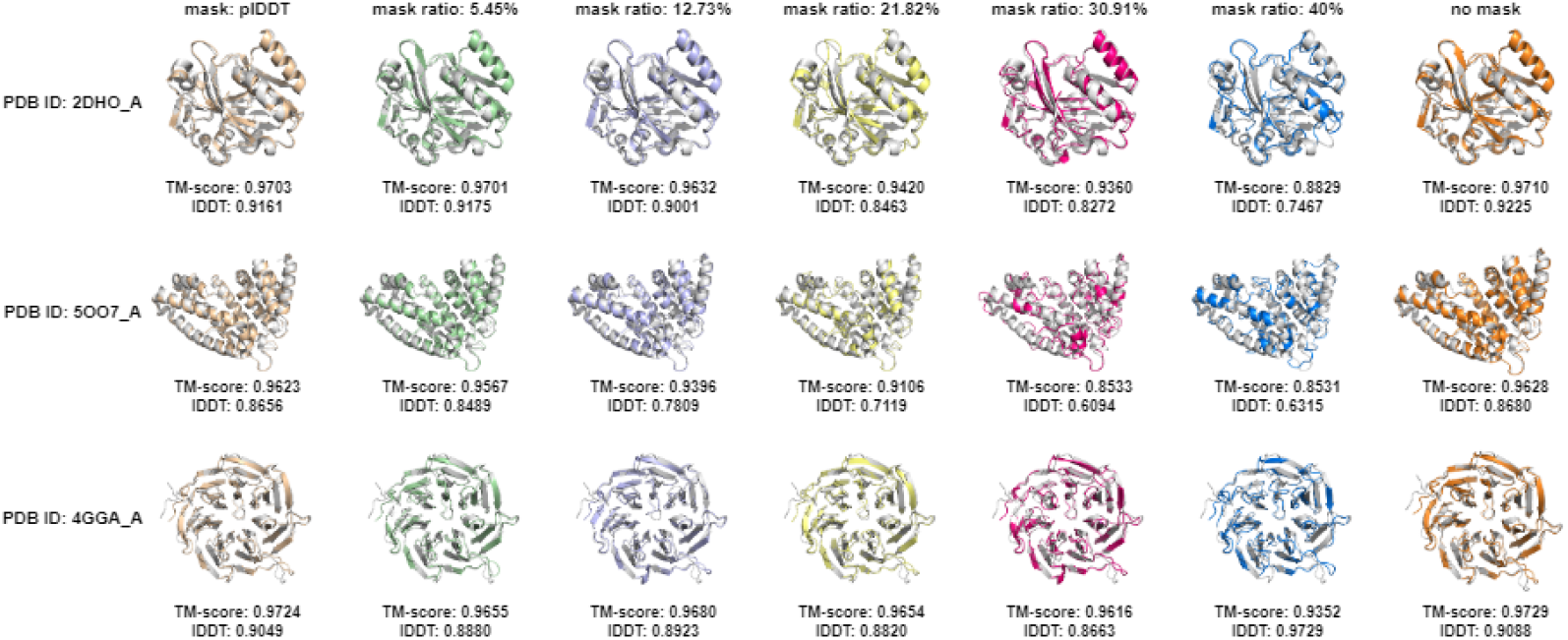
Decoy structures generated using geometric information predicted by SPIRED.

**Figure S5.**
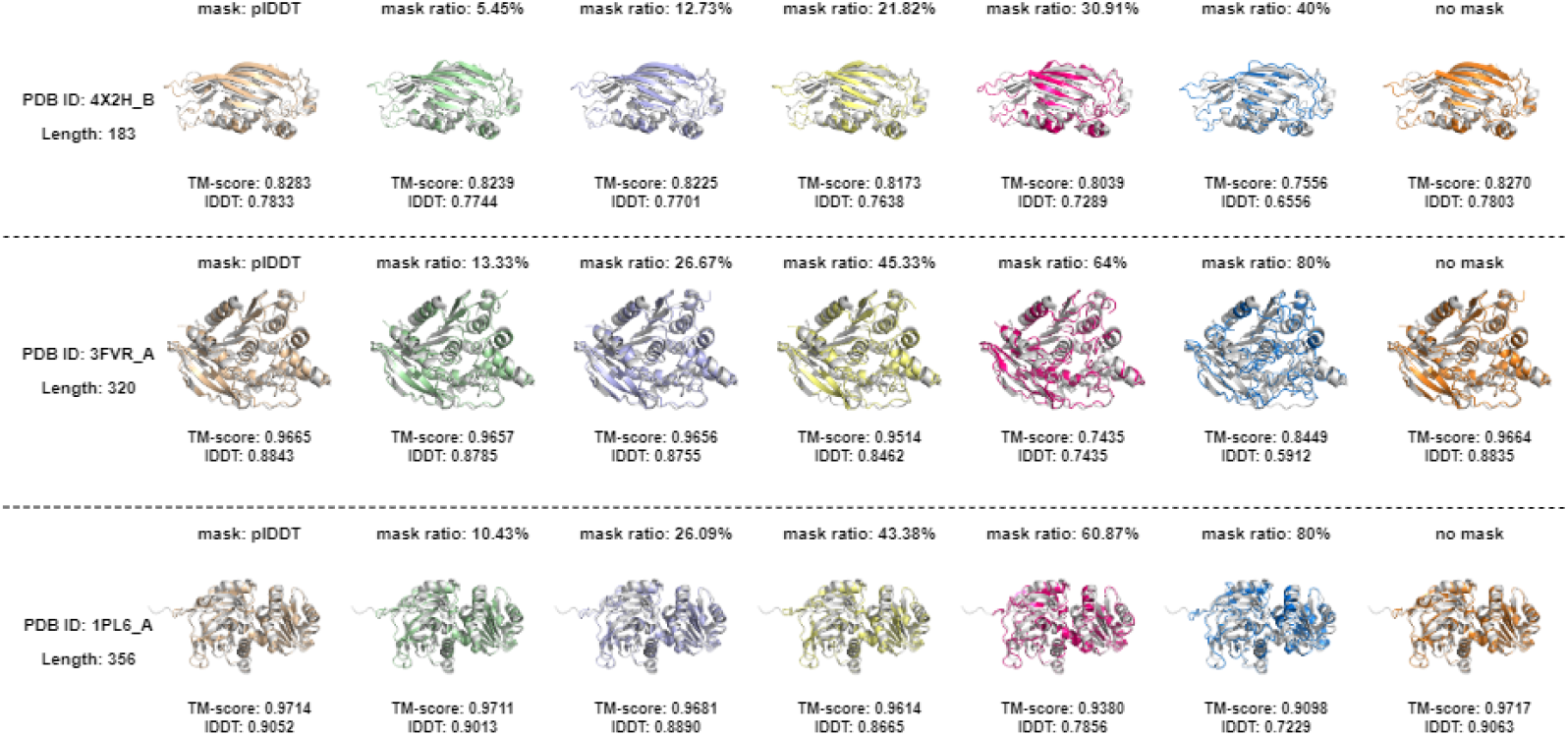
Decoy structures generated using geometric information predicted by Cerebra.

As described in **Methods**, the decoy structures are obtained by multiplying various mask weights to predicted geometric information and converting the masked 2D information into 3D structures through GDFold2 in parallel. The quality of these structures decreases with the increase of the mask ratio. As shown in Figures S4 and S5, geometric information predicted from SPIRED is more sensitive to masking than that from Cerebra. Eventually, the upper limit of mask ratio is set as 40% for SPIRED and 80% for Cerebra (see Figure 5A).

#### 2.2 QA model performs well in selecting the best prediction result

**Figure S6.**
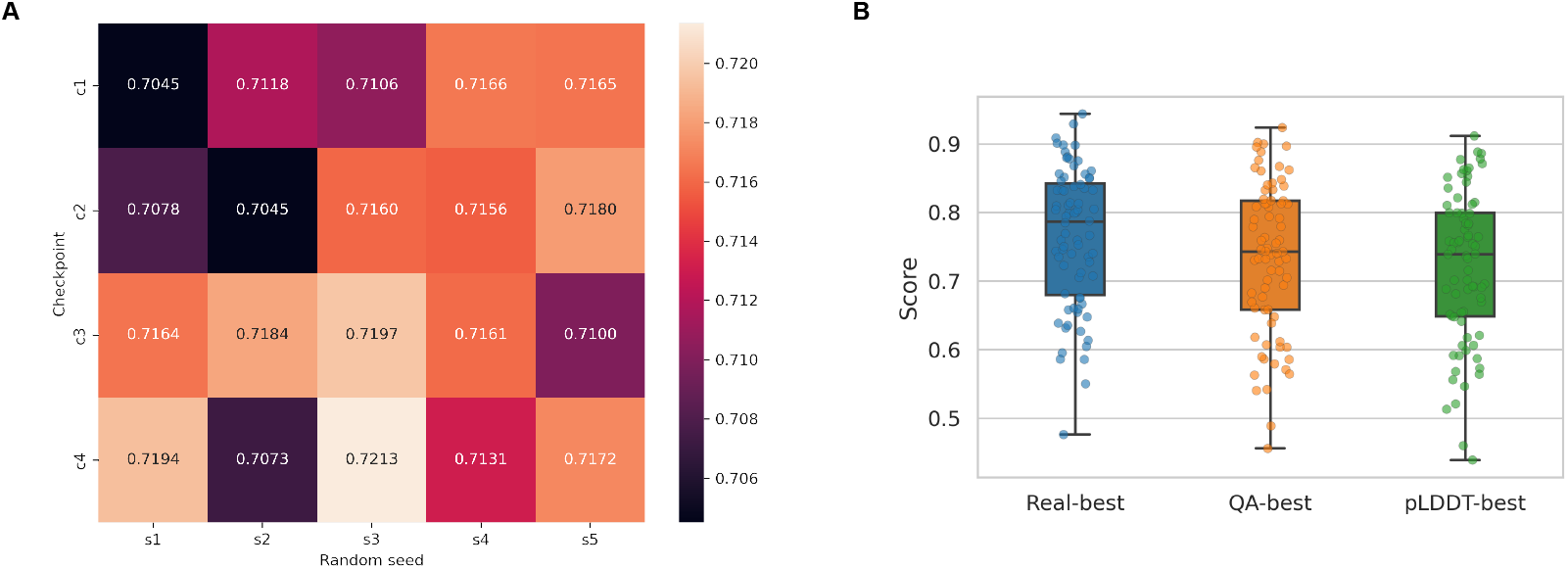
True evaluation score distribution of the test dataset. **A)** Average true evaluation scores of the testing dataset. **B)** Comparison of the best structures selected by the QA model and by plDDT reported by Cerebra.

Figure S6A summarizes the average true evaluation scores of the 76 targets in the CAMEO testing dataset folded by GDFold2 using geometric information predicted by 20 Cerebra models of different random seeds and checkpoints. We utilized the QA model to select the best one among the 20 structural models generated for each target protein and compared the entire results with those selected by plDDT. As shown in Figure S6B, the best structural models selected by the QA model have an average score of 0.7343, closer to the truly optimal structures (average score of 0.7643) than those picked up by the raw plDDT values reported by Cerebra (average score of 0.7209).

**Figure S7.**
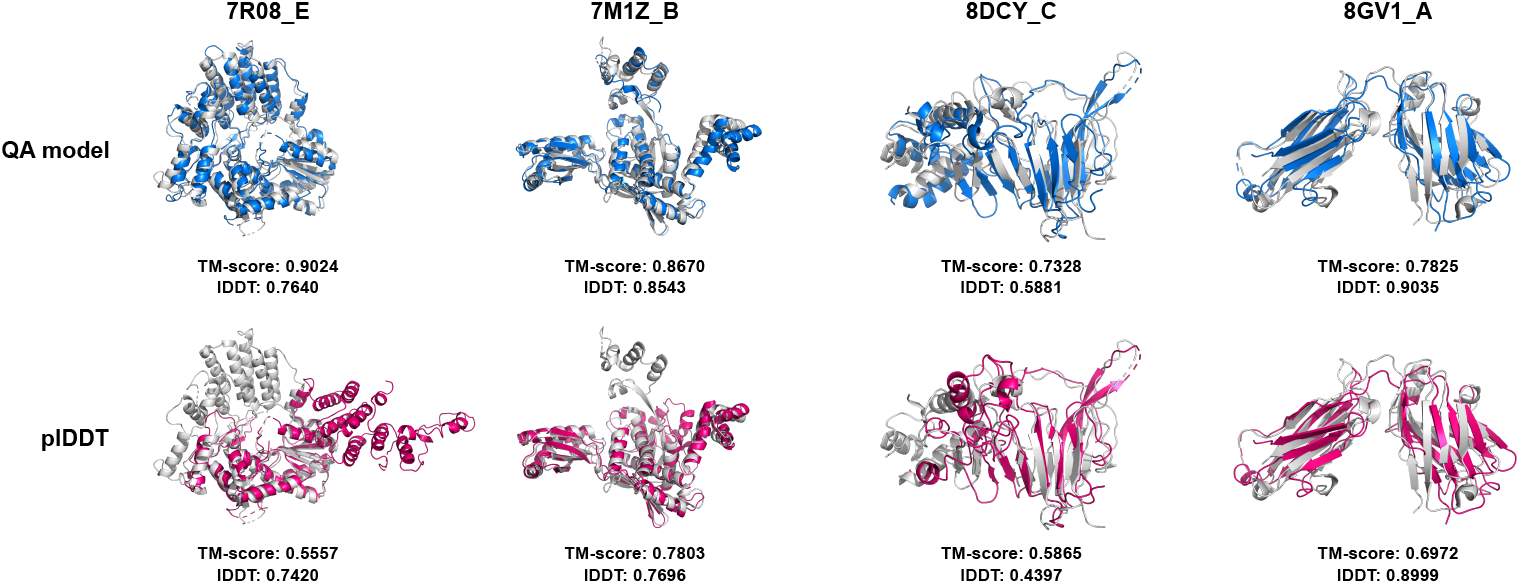
Comparison of the selected best structures.

Based on Figure S6 and showcases in Figure S7, the structures chosen by the QA model have higher quality than those selected by the plDDT reported by SPIRED/Cerebra, in terms of TM-score, lDDT and the rationality upon visual inspection.

### 3 Supplementary Details for Structural Dynamics

#### 3.1 Exploring the structural dynamics of ADK using AlphaFold2-predicted structures

**Figure S8.**
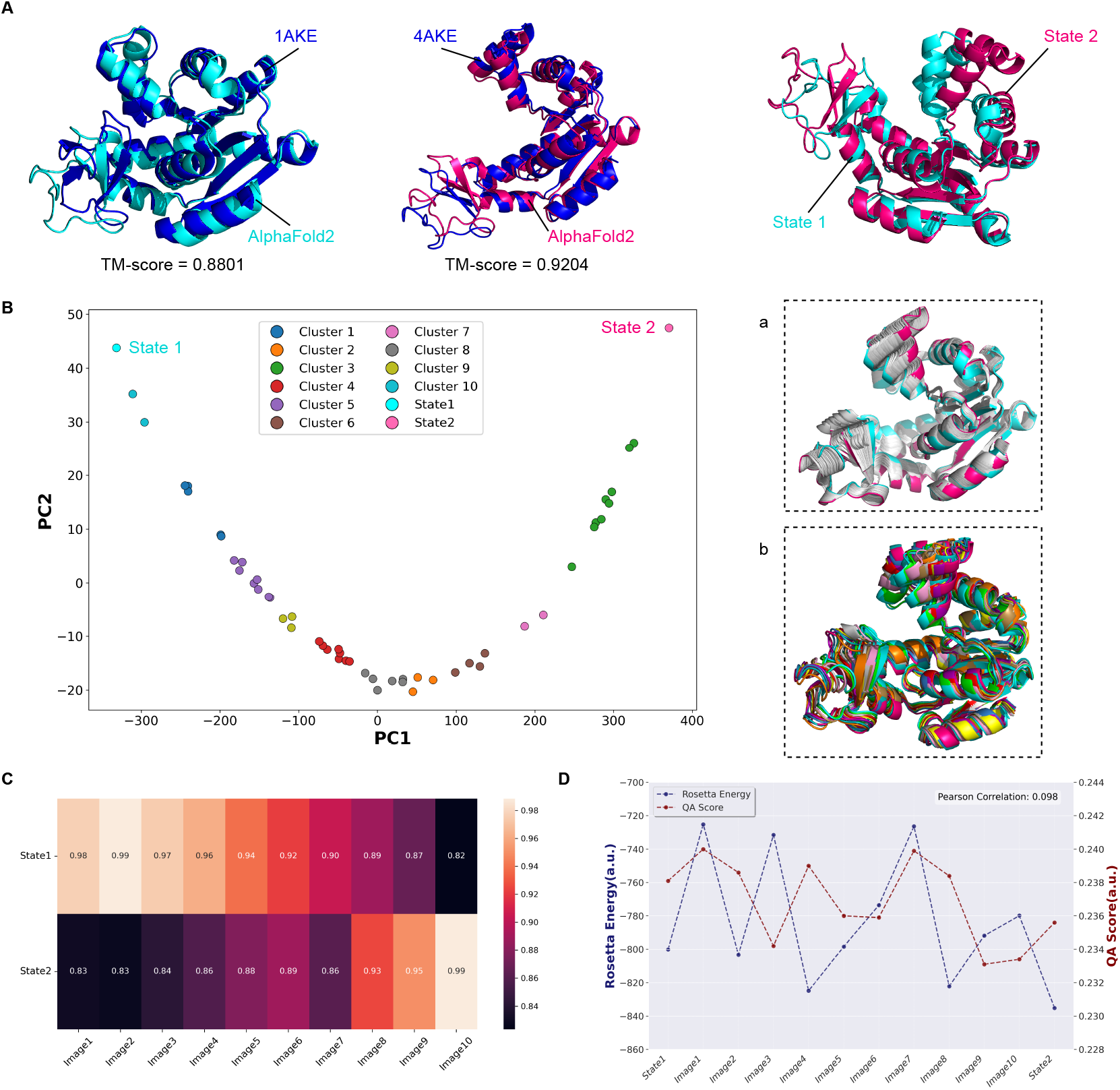
Transition path prediction using the multiple conformations predicted by AlphaFold2. This figure complements Figure 6 by using AlphaFold2-predicted structures as the substitutes for the experimental structures. **A)** The two conformational states of ADK could be successfully predicted by AlphaFold2 by carefully choosing the sequences from the MSA as input. **B-D)** Analogous counterparts to Figure 6C-E.

Following the procedure summarized in Figure 6A, the transition path between the two states of ADK could be constructed in a similar approach, except that the two structural states are predicted from AlphaFold2^3^ by MSA manipulation. As shown in Figure S8A, the AlphaFold2-predicted structure models resemble the experimental structures of 1AKE and 4AKE, respectively. Hence, the transition path could be constructed in Figure S8B-D, like Figure 6C-E.

### 4 Algorithm and Pseudocode for GDFold2 and the QA model

#### Algorithm 1 Constraints based on vectors predicted by Cerebra

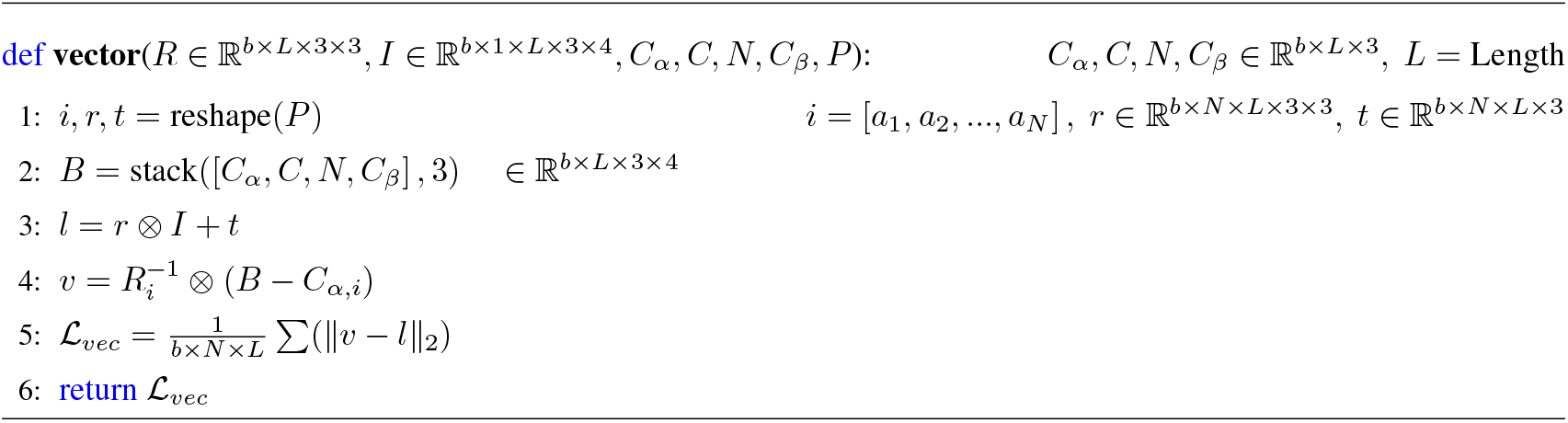

#### Algorithm 2 Constraints based on vectors predicted by SPIRED

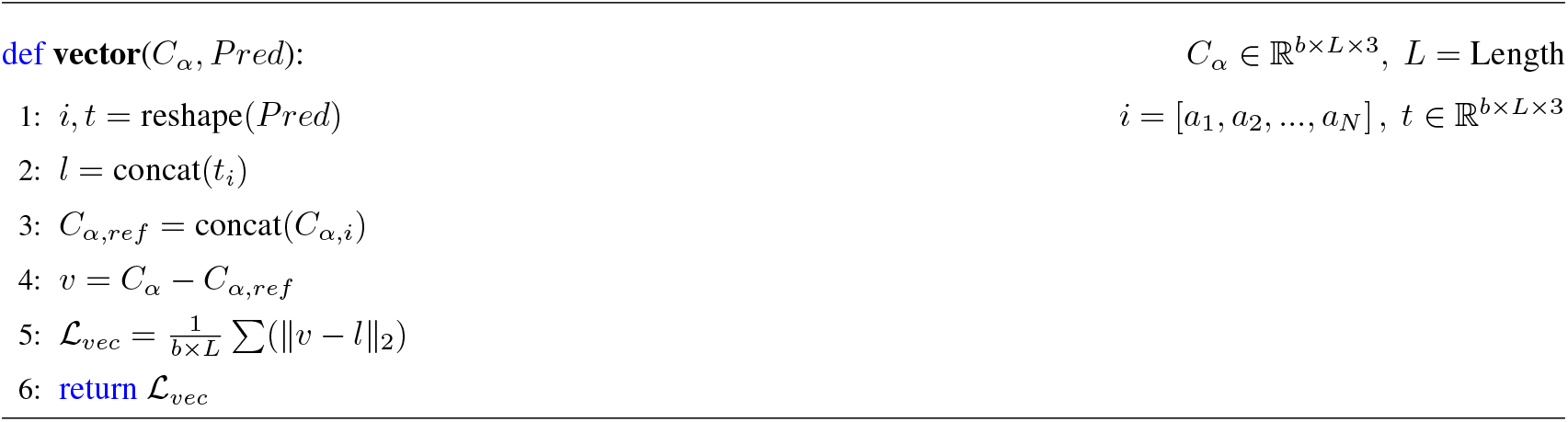

#### Algorithm 3 Infer the internal coordinate of the previous *C* atom in the local coordinate system of the current residue

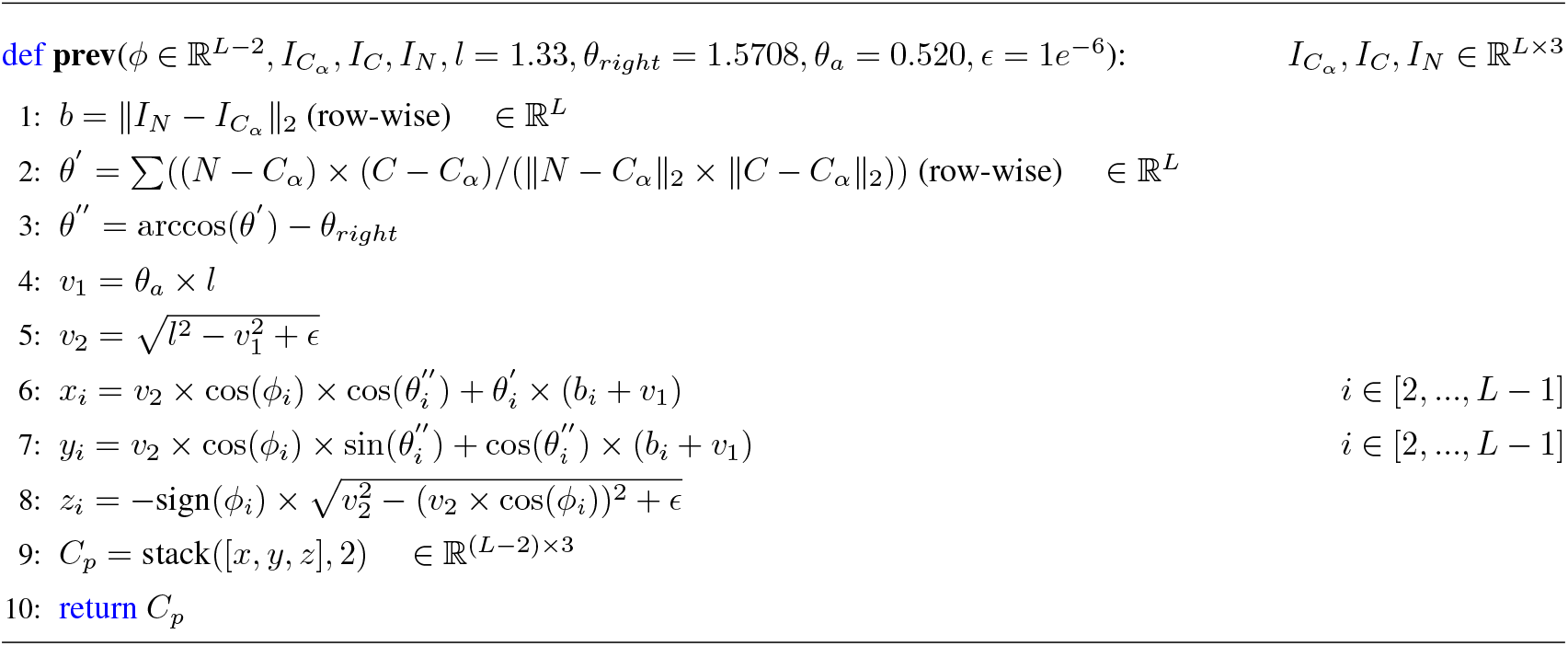

#### Algorithm 4 Infer the internal coordinate of the next *N* atom in the local coordinate system of the current residue

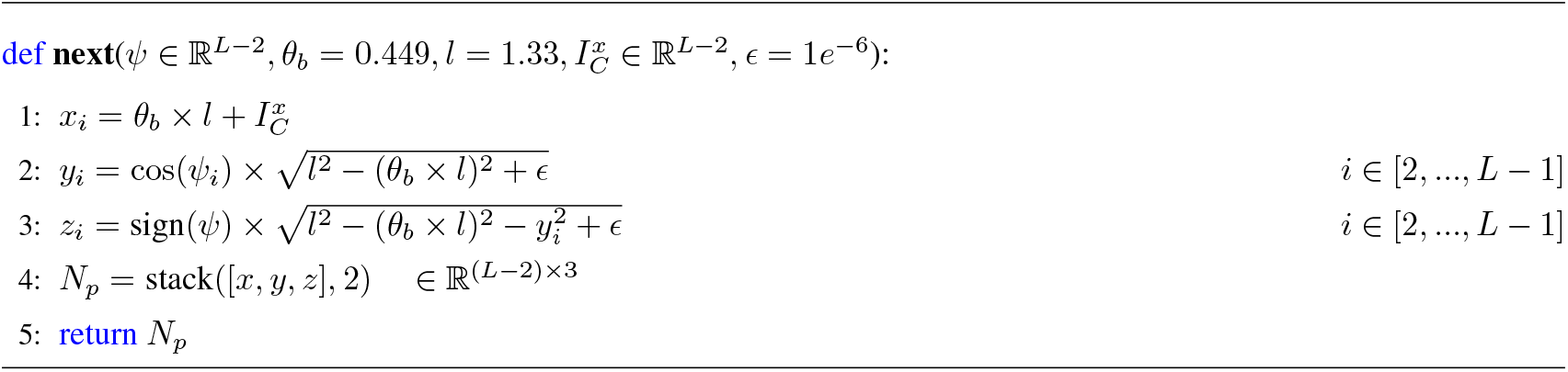

#### Algorithm 5 Constraints on the positions of preceding *C* and next *N* atoms based on predicted dihedral angles

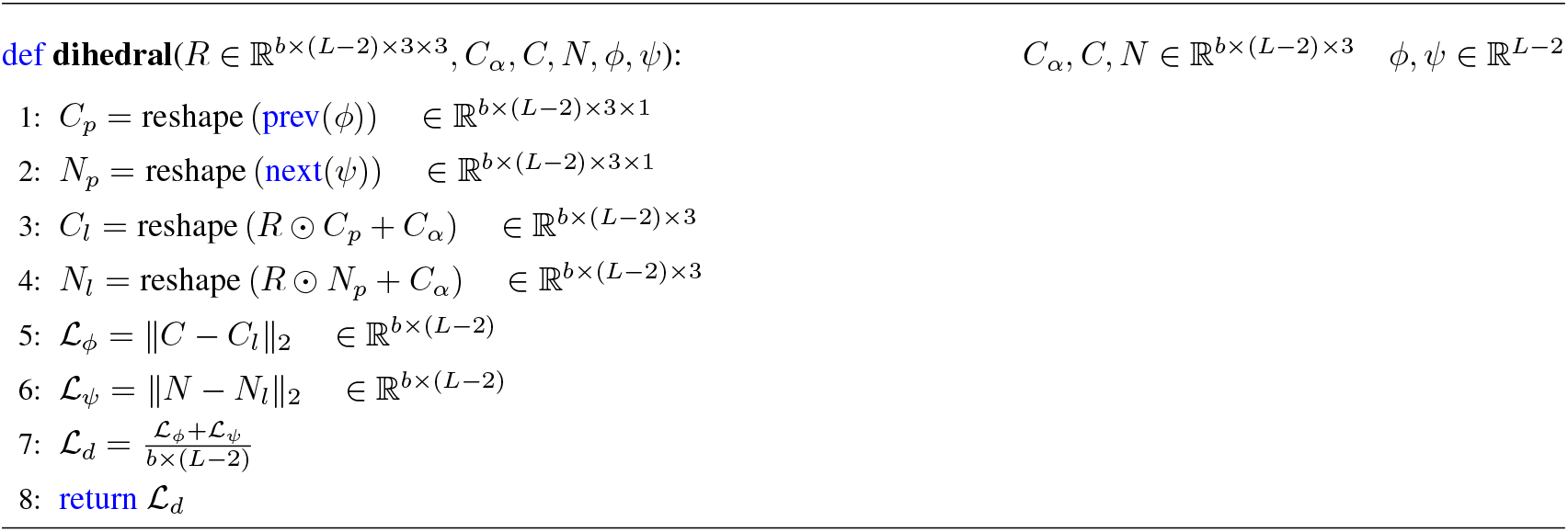

#### Algorithm 6 Peptide plane constraints

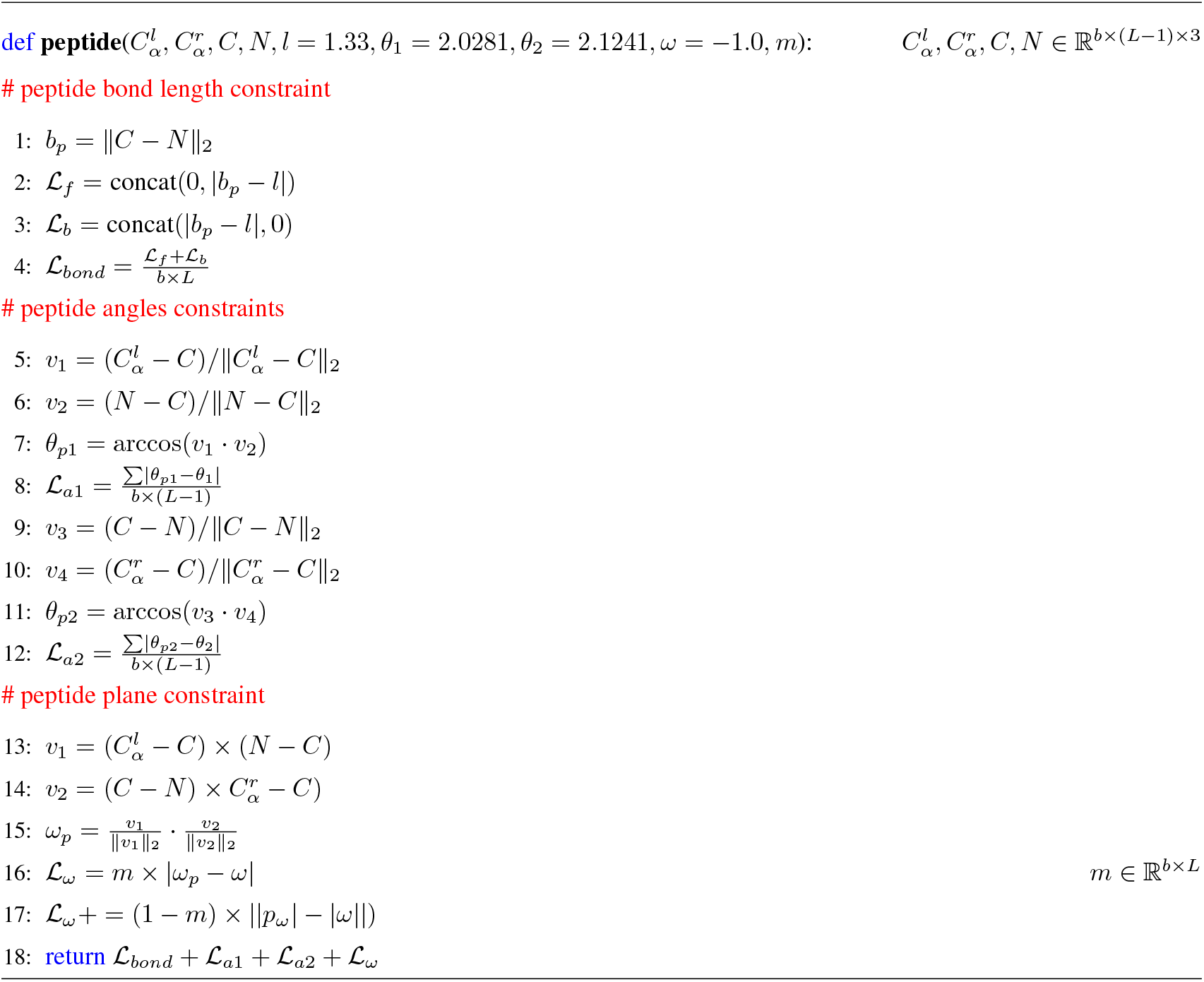

#### Algorithm 7 *C*_*α*_-*C*_*α*_ distance constraint

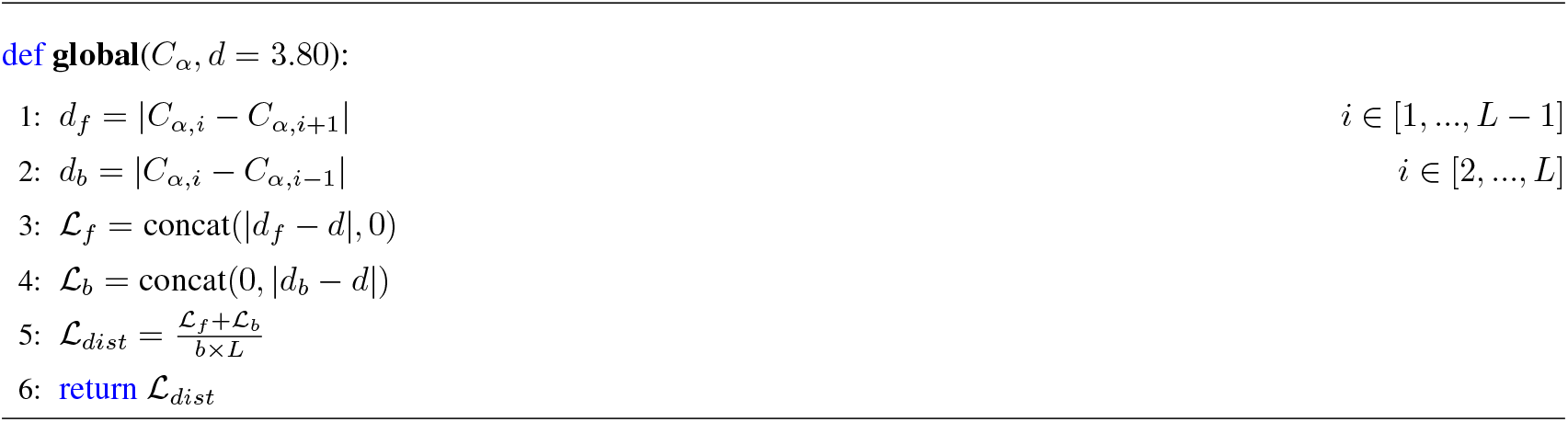

#### Algorithm 8 Van der Waals repulsion constraint

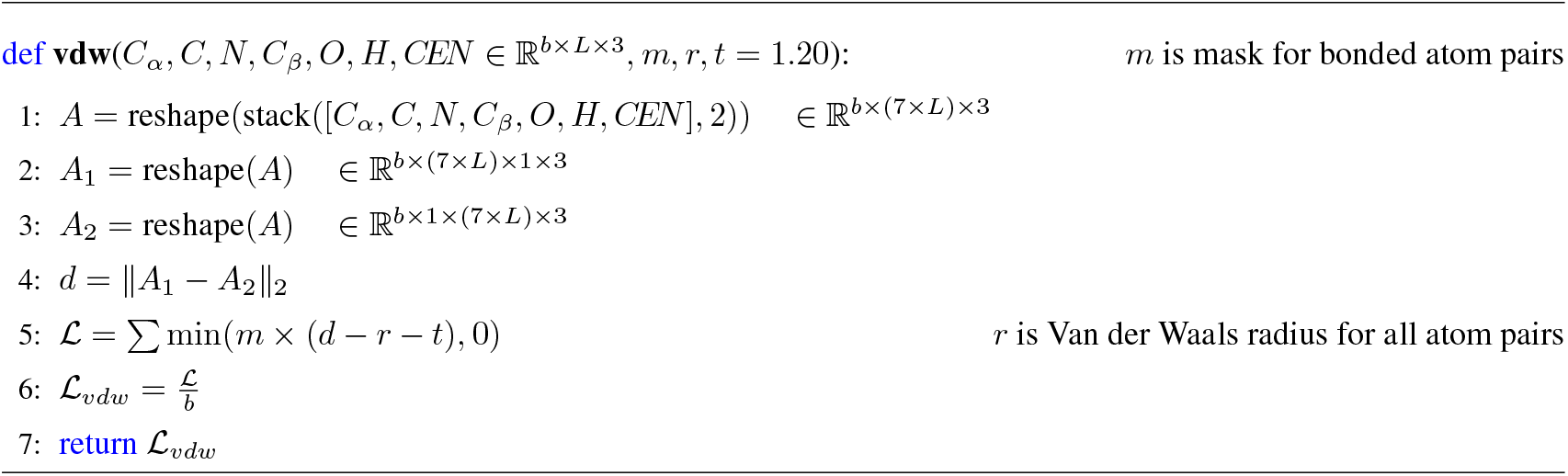

#### Algorithm 9 Convert Euler Angle into rotation matrix

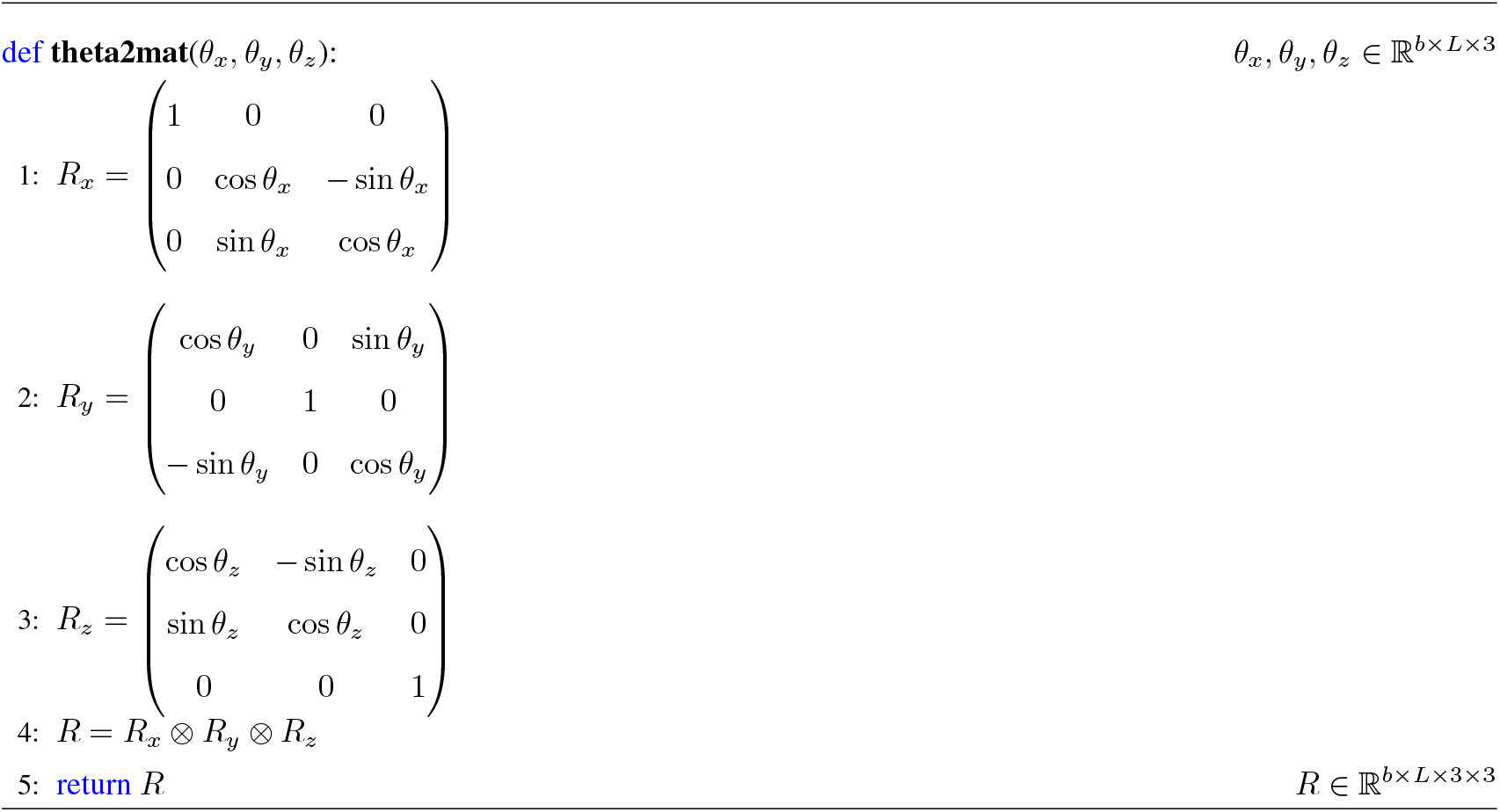

#### Algorithm 10 Calculate the rotation matrix of the local coordinate system of a residue

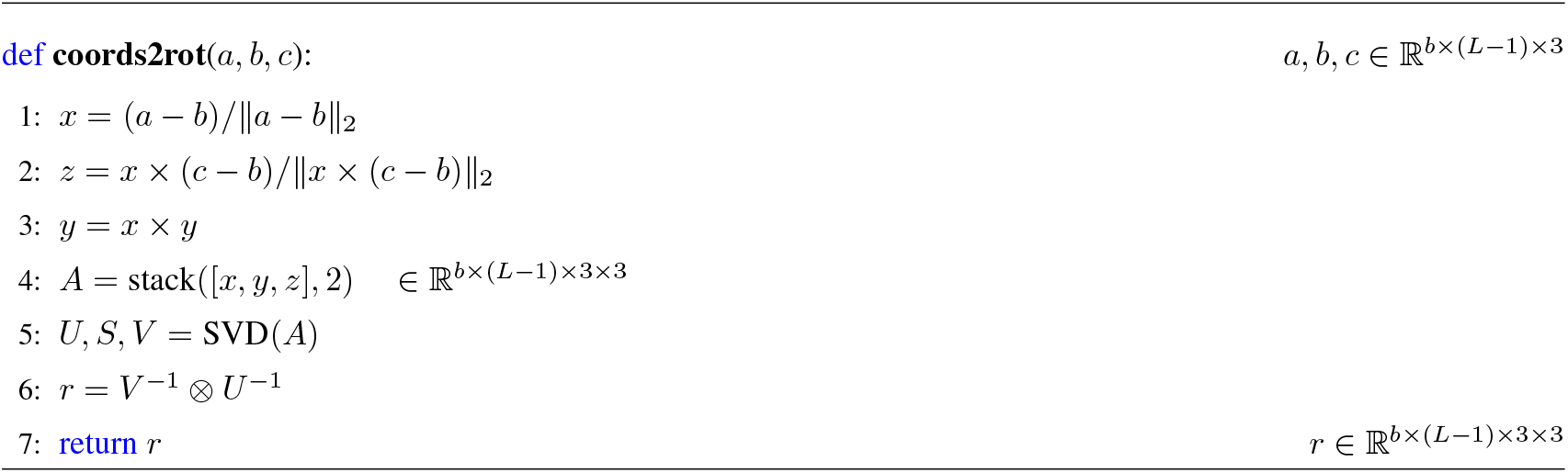

#### Algorithm 11 Infer the global coordinates of oxygen atoms

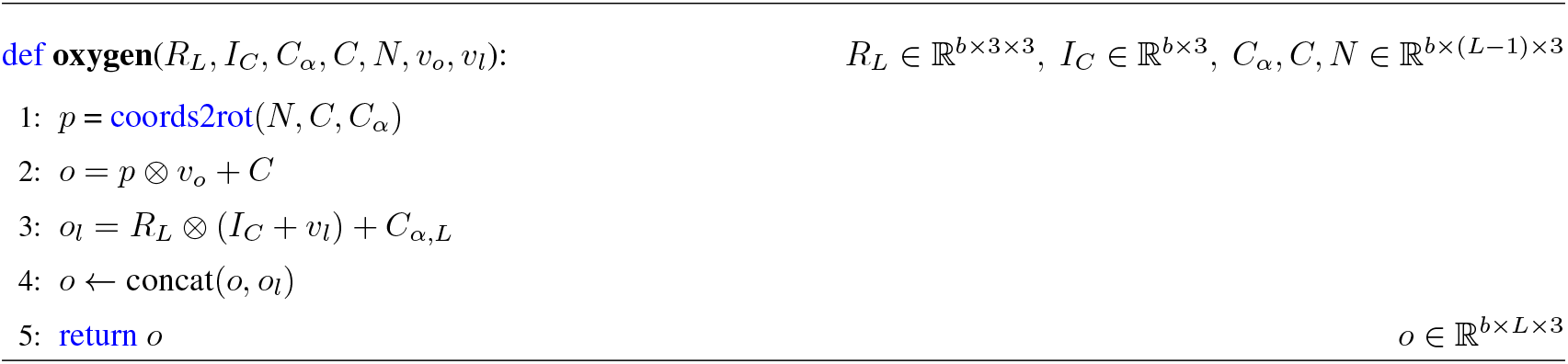

#### Algorithm 12 Infer the global coordinates of hydrogen atoms

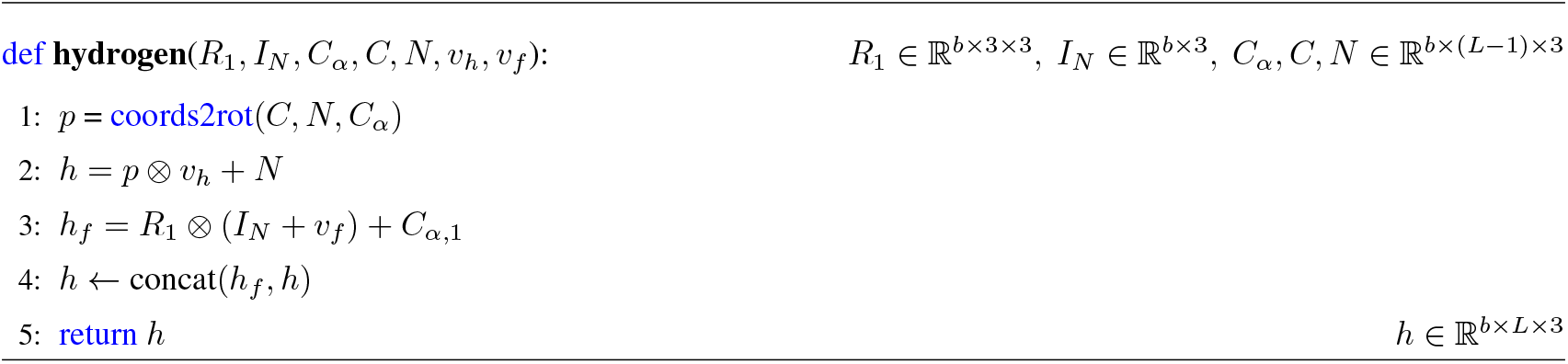

#### Algorithm 13 Calculate the coordinates of backbone atoms and the rotations of residue coordinate systems

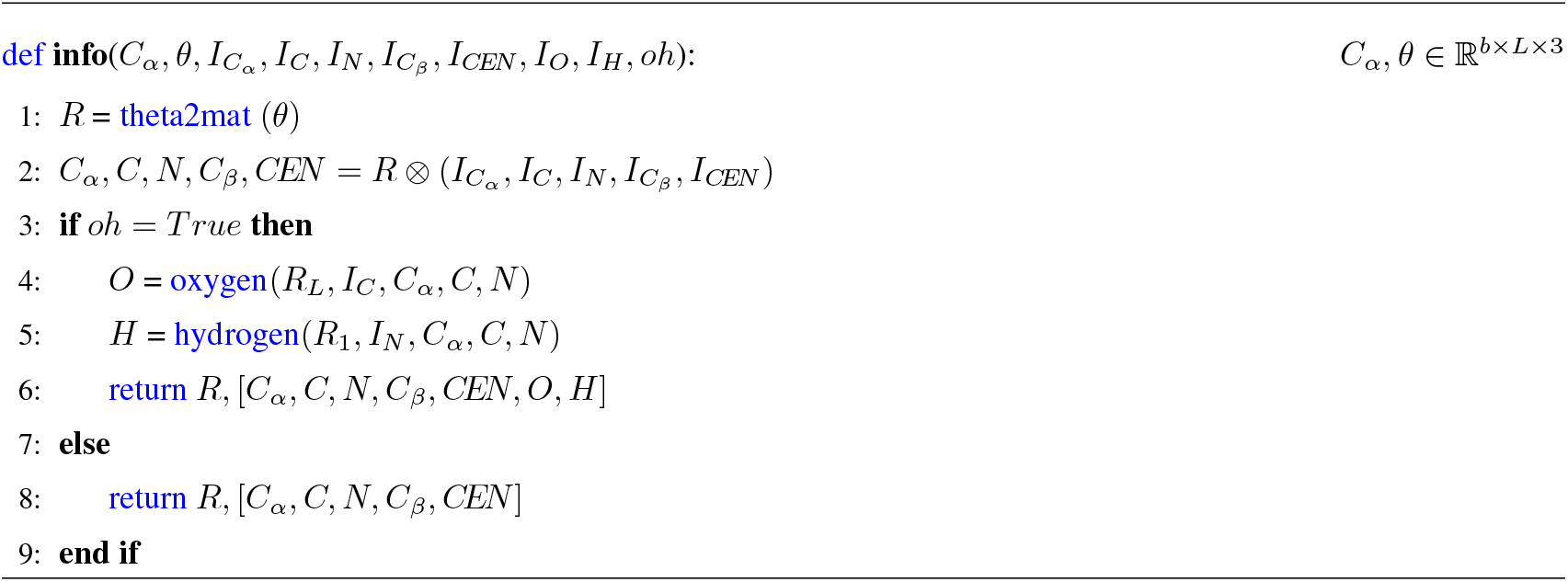

#### Algorithm 14 Protein folding process

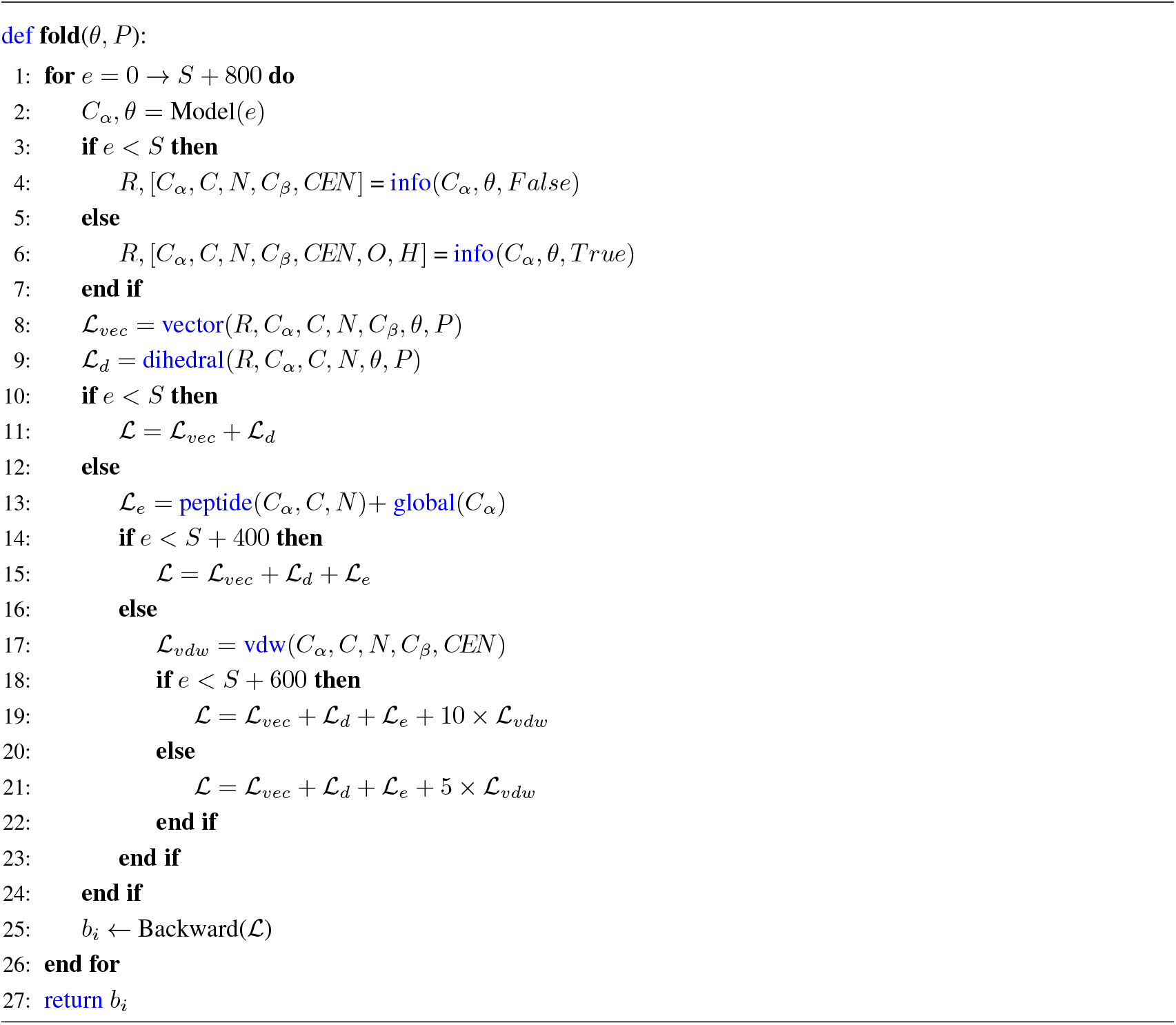

#### Algorithm 15 Graph Vector Perceptron (GVP)

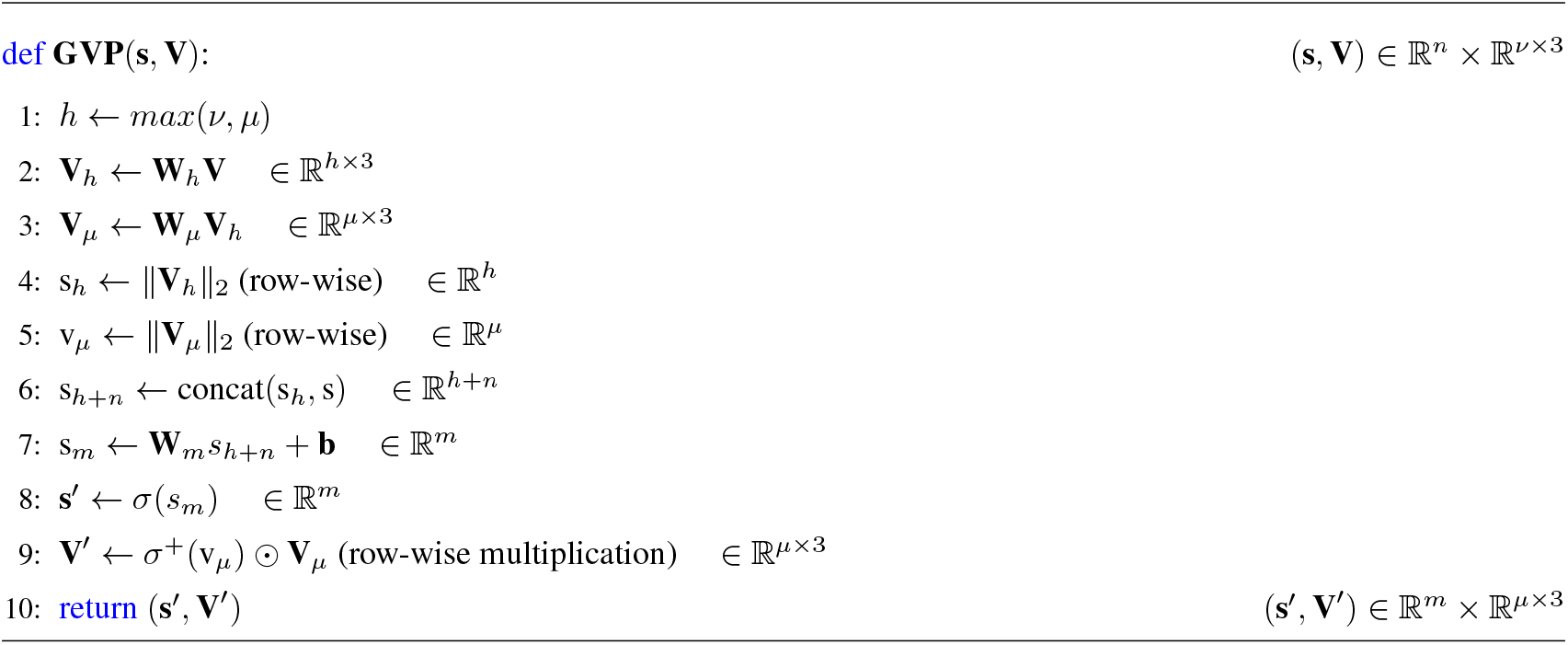

#### Algorithm 16 Masked geometric constraints for structural dynamics

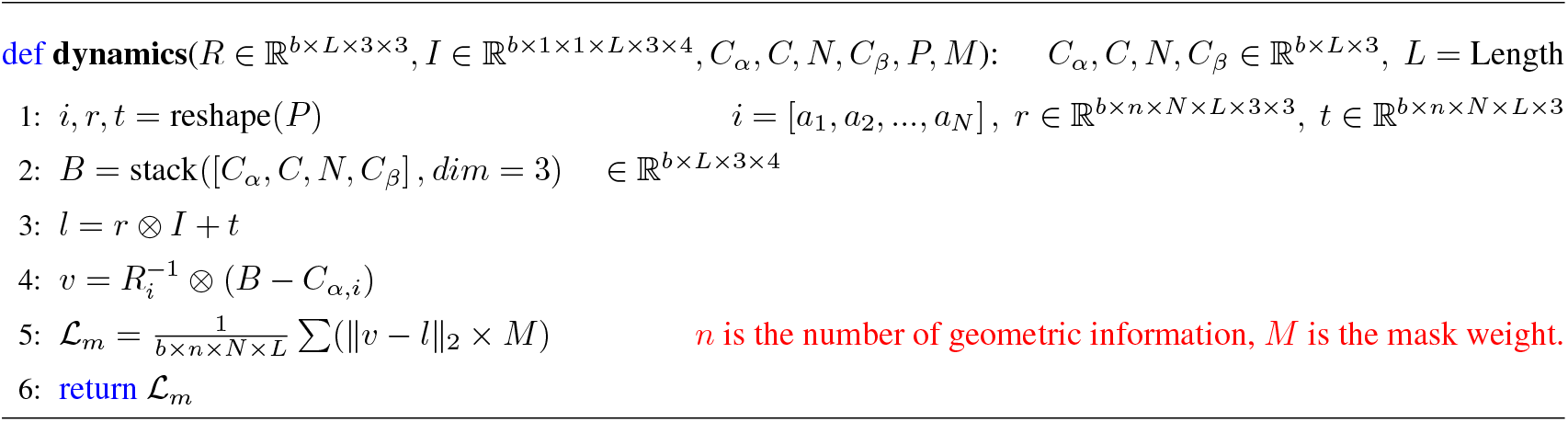

## Notes

### Competing Interest Statement

The authors have declared no competing interest.

### Summary of Updates

Authors updated; Sever link added in Summary; Investigation of protein conformational dynamics added in Methods, Results, and Supplementary Information.

